# Pioneer bark beetle attacks induce multifaceted localized defense responses in Norway spruce

**DOI:** 10.1101/2025.08.06.668443

**Authors:** Marcelo Ramires, Sigrid Netherer, Martin Schebeck, Karin Hummel, Sarah Schlosser, Ebrahim Razzazi-Fazeli, Reinhard Ertl, Muhammad Ahmad, Ana Espinosa-Ruiz, Esther Carrera, Erwann Arc, Maria Ángeles Martínez-Godoy, Jorge Baños, Teresa Caballero, Thomas Ledermann, Marcela van Loo, Carlos Trujillo-Moya

## Abstract

Conifer forests worldwide are increasingly threatened by environmental stressors, especially bark beetle infestations. In a novel experimental field setup with controlled attacks on clonal trees, we investigated the molecular defense responses of 35-year-old Norway spruce trees to colonization attempts by the spruce bark beetle, *Ips typographus*. A multi-omics approach was used to study tree responses to bark beetle pioneers under natural conditions. At the site of the attacks, increased levels in jasmonic acid, together with changes in other phytohormones, induced differential gene expression of up to 1,900 genes from multiple defense-related pathways. These transcriptomic alternations were corroborated by proteomic and metabolomic shifts, which ultimately showed increased levels of compounds involved in deterring bark beetle attacks, such as phenolic aglycones, diterpene resin acids, mono- and sesquiterpenes, and lignin. Our study highlights the intricate yet rapid and effective local defense response of Norway spruce against pioneer bark beetle attacks and lays a foundation for in-depth studies of this and other conifers resistance to phloem-feeding insects.

## Introduction

Forests worldwide provide essential ecosystem services, including carbon and water cycling, biodiversity maintenance, and timber production. Norway spruce (*Picea abies* (L.) H. Karst.), a dominant species in boreal and temperate Eurasian forests, is especially important in subalpine and montane regions. In these landscapes, natural spruce stands play a vital role in stabilizing soil, regulating water resources, and protecting infrastructure and settlements from natural hazards, such as avalanches and landslides (Smith et al., 2005). The widespread distribution of Norway spruce under diverse environmental conditions highlights its ecological and economic importance (Chakraborty et al., 2021, 2024; Mauri et al., 2022; “The State of the World’s Forests 2020”, 2020).

Triggered by climate change, Norway spruce, like other conifers, is increasingly exposed to abiotic stressors such as heat, drought, and extreme weather events like storms. The severity and frequency of these disturbances have been intensifying and are expected to increase in the future, potentially altering the dynamics of tree susceptibility and resistance to insect infestations (Allen et al., 2010; Dyderski et al., 2018; Jactel et al., 2012; Rouault et al., 2006). In recent decades, the Eurasian spruce bark beetle, *Ips typographus* (L.) (Coleoptera: Curculionidae: Scolytinae), has caused extensive damage to Norway spruce forests during epidemic population outbreaks, often initiated by preceding abiotic disturbances (Bentz et al., 2019; Hlásny et al., 2021). Bark beetle infestations, documented for centuries in managed forests of Europe (Lehmanski et al., 2023; Marini et al., 2017), not only result in significant timber losses, but also in broader impacts on ecosystem services (Raffa et al., 2015; Seidl et al., 2015). The recent increased frequency and severity of these outbreaks highlights the urgent need to understand and mitigate the impacts of bark beetle attacks on forest ecosystems (Allen et al., 2010; Dyderski et al., 2018). While bark beetle mass attacks inevitably lead to extensive stand mortality, constitutive and induced chemical defenses can effectively protect spruce trees against the first invaders, so-called pioneer beetles (Netherer et al., 2021; Schiebe et al., 2019).

Extensive research has investigated the constitutive and inducible anatomical and biochemical defense strategies of conifers against biotic invaders, including Norway spruce and its interactions with *I. typographus*. Key defense mechanisms include radial and axial resin duct formation, as well as metabolite storage and synthesis in epithelial or polyphenolic parenchyma cells (Franceschi et al., 2005; Krokene, 2015; Krokene et al., 2008; Netherer & Hammerbacher, 2021). Similar defensive strategies have been extensively studied in other conifer species, such as lodgepole pine (*Pinus contorta*) and its interactions with the mountain pine beetle (*Dendroctonus ponderosae*), providing broader insights into conifer resistance to phloem-feeding insects (Clark et al., 2012; Howe et al., 2024; Raffa & Berryman, 1982). The defense strategies of conifers against insect herbivores and pathogens are tightly connected to specialized biosynthetic pathways that produce a diverse array of secondary metabolites. Among these, terpene biosynthesis pathway plays a crucial role in conifer defense, generating a wide range of volatile (mono- and sesquiterpenes) and non-volatile (diterpenes) compounds. These substances are involved in both direct toxicity and signaling in a dose-dependent way (Erbilgin et al., 2007; Keeling & Bohlmann, 2006; Miller & Borden, 2000), and are main constituents of resin, a strong physical and chemical barrier against bark beetles and their fungal associates (Martin et al., 2004). Additionally, the phenylpropanoid, flavonoid and stilbene biosynthesis pathways lead to the production of compounds that can interfere with insect feeding and digestion (Hammerbacher et al., 2011, 2019; Treutter, 2006). Given the long-standing co-evolutionary history between *I. typographus* and conifers of the Pinaceae family (Cognato, 2015), these secondary metabolite biosynthetic pathways likely represent a conserved yet complex immune response that warrants further exploration.

Natural outbreaks, while valuable for retrospective molecular analyses, present inherent technical constraints for studying immediate tree defense reactions to initial bark beetle attacks. The unpredictability and broad spatial scale of these infestations make it challenging to control experimental conditions. To account for attack density and timing, field (no)-choice bioassays have been employed in conjunction with physiological and biochemical investigations of trees from different genotypes (Basile et al., 2024; Netherer et al., 2024). While many studies aiming to elucidate the molecular defense mechanisms of Norway spruce have relied on the artificial application of methyl jasmonate (MeJA) or inoculations with bark beetle-associated fungi, our study utilizes direct bark beetle attacks under controlled natural conditions. This approach provides a more ecologically realistic and integrative perspective on tree defense responses (Erbilgin et al., 2006; Wilkinson et al., 2022; Zhao et al., 2010). We applied a comprehensive experimental approach involving controlled bark beetle attacks within a clonal stand of 35-year-old Norway spruce trees. This unique setup minimized genetic variability and enabled us to investigate the complete molecular defense response to colonization attempts by *I. typographus*. Our approach provides a detailed, highly time-resolved characterization of the dynamic molecular defense response in Norway spruce during the critical early phase of colonization by pioneer bark beetles, spanning up to seven days post-exposure. By integrating transcriptomic, proteomic, hormonal, and metabolite profiling analyses, this multi-omics framework offers unprecedented insights into the complex, localized defense strategies of Norway spruce deployed at the onset of attack. This work establishes a foundation for further investigations into conifer resistance mechanisms under biotic stress, with broader implications for understanding tree resistance to insect infestations.

## Results

### Bark beetle attacked trees during a controlled exposure with individual cages

To investigate the initial defense responses of Norway spruce to bark beetle colonization, we exposed 20 clonal, 35-year-old Norway spruce trees to controlled attacks by male beetles (i.e., the gallery-establishing sex). Additionally, 20 clonal trees were used as negative control. All trees in the attack group (Figure 1A) were affected by bark beetles, each tree being attacked by at least three out of a maximum of six beetles. We considered any beetle damage to the trees as an attack, ranging from surface scratches on the bark (i.e. feeding/boring attempts) to burrowing into the outer bark/phloem. On the second day after exposure (i.e. first observation), beetle activity was evidenced by signs of early colonization (scratches on the bark surface). By day three after exposure, we observed that some beetles had burrowed into the outer bark and phloem to form mating chambers (i.e., start of gallery establishment), while others remained on the surface, feeding on the outer bark. By the end of the assay (day 7), beetles that penetrated the phloem were consistently found dead and encased in resin. No beetles other than those used for our controlled attacks were present on the attacked trees, and negative control trees remained entirely free of any signs of beetle infestations (Figure 1B). In attacked trees, bark/phloem samples were taken from the colonization site to analyze the local defense response (LR) and from the opposite side of the trunk to assess the systemic defense response (SR). Additionally, samples from negative control trees (NC) were collected to distinguish beetle-induced responses from background physiological variation.

**Fig. 1.**
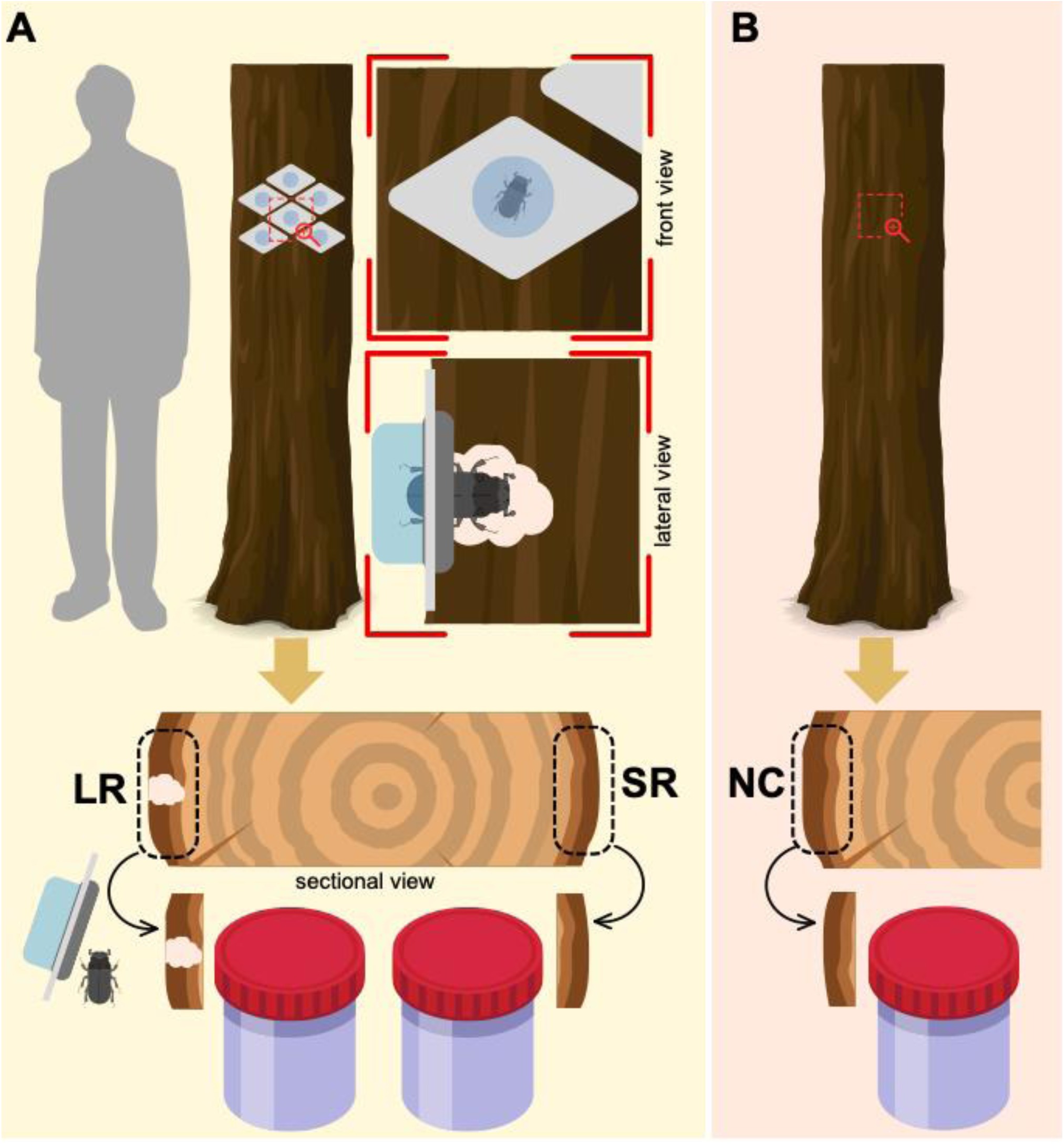
Schematic representation of the experimental design and sampling. **A** Setup for bark beetle exposure on the attacked trees. Beetles were introduced into individual cages (i.e. capsular pits) attached to the surface of the tree trunk by rubber bands. Local response (LR) samples were collected from the site of beetle attack, while systemic response (SR) samples were collected from the diametrically opposite side of the trunk. **B** Negative control (NC) trees were not exposed to beetles and were sampled at the same height and orientation as in the LR group for molecular comparison.

### RT-qPCR revealed a strong and dynamic local defense response

To assess differences in gene expression patterns associated with defense responses, we performed a Principal Component Analysis (PCA) of 19 biotic stress-related genes (Supplementary Table 1 and 2). These included key genes involved in signal transduction and environmental adaptation pathways, such as plant-pathogen interactions (e.g., Calmodulin), MAPK signaling (e.g., MPK6), and plant hormone signal transduction (e.g., MYC2), as well as in the biosynthesis of plant secondary metabolites like phenylpropanoids (e.g., HCT), flavonoids (e.g., FH3), and terpenoids (e.g., FPPS). The analysis revealed a clear distinction between the attacked and control trees for the local response, but not for the systemic response (Figure 2A) (Supplementary Figure 1). In particular, LR samples showed a significant up-regulation of most genes as early as two days after the attack (e.g., MA_10435304g0010; Flavanone-3-Hydrolase [F3H]) (Figure 2B), whereas SR samples did not display any significant changes compared to the negative control for the majority of genes. In addition, we analyzed the expression of multiple pDIR family genes using degenerated primers designed to amplify 10 dirigent protein-coding genes (Supplementary Figure 2). This analysis revealed a strong increase in expression within the LR group at all time points, with a peak fold change of 46.45 on day 5 (Figure 2C). These genes promote the stereoselective formation of lignans (i.e., water-soluble antifeedant polyphenols smaller than lignin). In contrast, the SR group only showed a ≈2-fold rise on day 3, while the other time points stayed close to baseline of the negative control. These results, when compared with the broader spectrum of responses from our pool of 19 analyzed genes, mirror the general gene expression patterns (Figure 2A), making this a promising defense expression marker. Since the SR samples showed minimal to no changes compared to the NC group throughout all analyzed time points, the subsequent analyses in this study focused exclusively on the comparison of LR and NC groups.

**Fig. 2.**
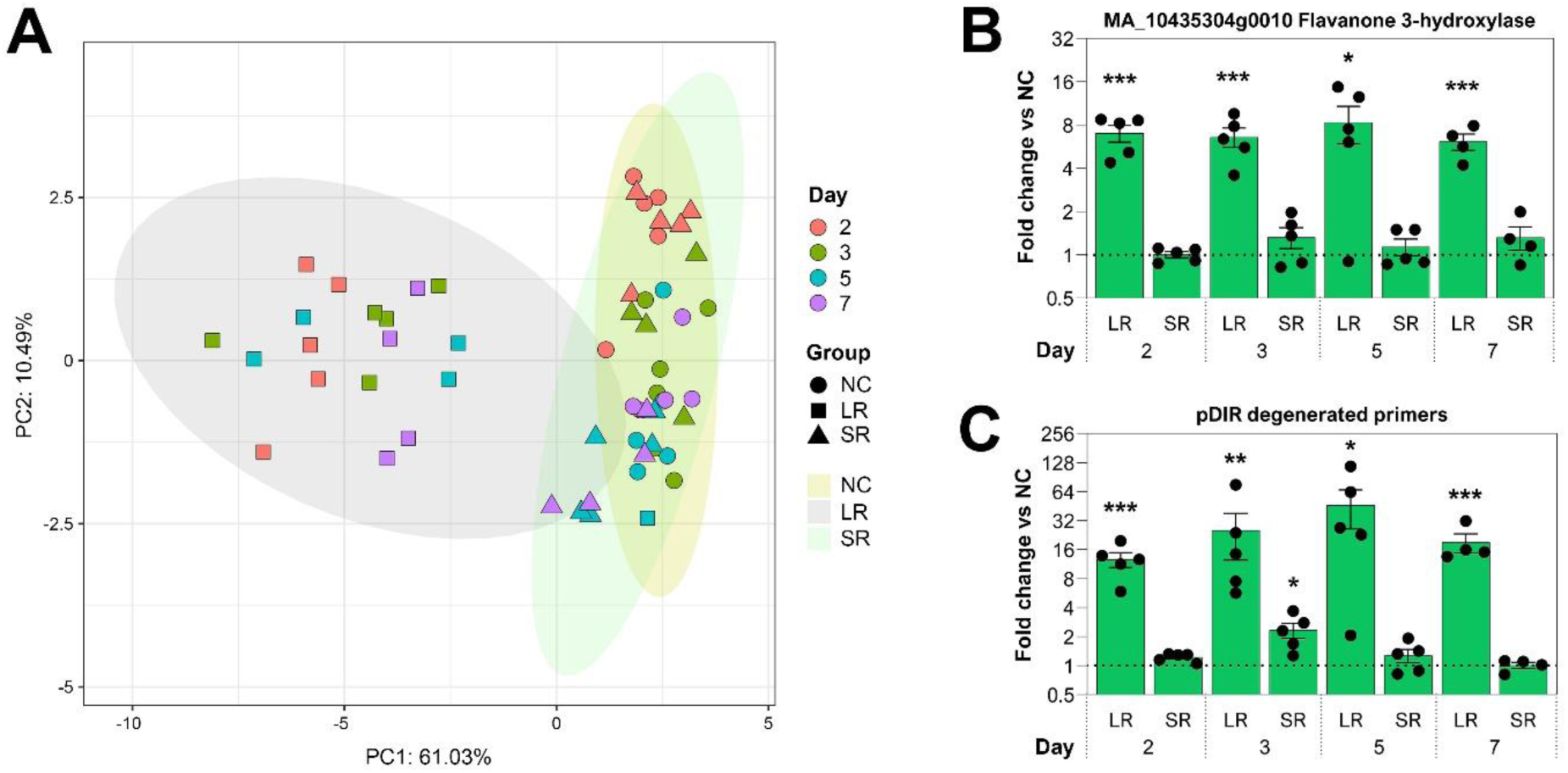
RT-qPCR analysis of differential gene expression in Norway spruce samples following a bark beetle attack. **A** Principal component analysis (PCA) of gene expression data showing a separation between negative control (NC), local response (LR), and systemic response (SR) at days 2, 3, 5, and 7 after bark beetle exposure. PC1 explains 61% of the variance, while PC2 explains 10% of the variance. **B** RT-qPCR results for the expression of the flavanone 3-hydroxylase (F3H) gene (MA_10435304g0010) in the LR and SR groups relative to the NC. **C** RT-qPCR results for the expression of PDIR gene family in the LR and SR groups relative to the NC. Statistical significance was determined using an unpaired *t*-test with Welch’s correction. Significance levels are indicated as follows: p < 0.05 (*), p < 0.01 (**), and p< 0.001 (***).

### Bark beetle attack locally boosts jasmonic acid and triggers multiple phytohormone shifts

Bark beetle attacks caused a significant increase in the major spruce defense signaling hormone jasmonic acid (JA) and a decrease in abscisic acid (ABA), indole-3-acetic acid (IAA), gibberellin A1 (GA1), certain cytokinins, and salicylic acid (SA) at the site of the attack (LR) (Figure 3A; Supplementary Table 3). Throughout the whole experimental period (day 2-7), JA levels rose considerably (up to 35-fold) in LR as compared to NC while ABA levels showed a marked decrease (up to 20-fold). IAA levels showed decreases on days 3 and 7, while GA1 decreased on days 5 and 7 (Figure 3A). Cytokinin levels fluctuated throughout, with isopentenyladenine (IP) decreasing on days 2 and 3, dihydrozeatin (DHZ) decreasing on day 7, and trans-zeatin (tZ) slightly increasing on day 5. Additionally, SA levels dropped only on day 5, while gibberellin A4 (GA4) remained stable throughout the observation period.

**Fig. 3.**
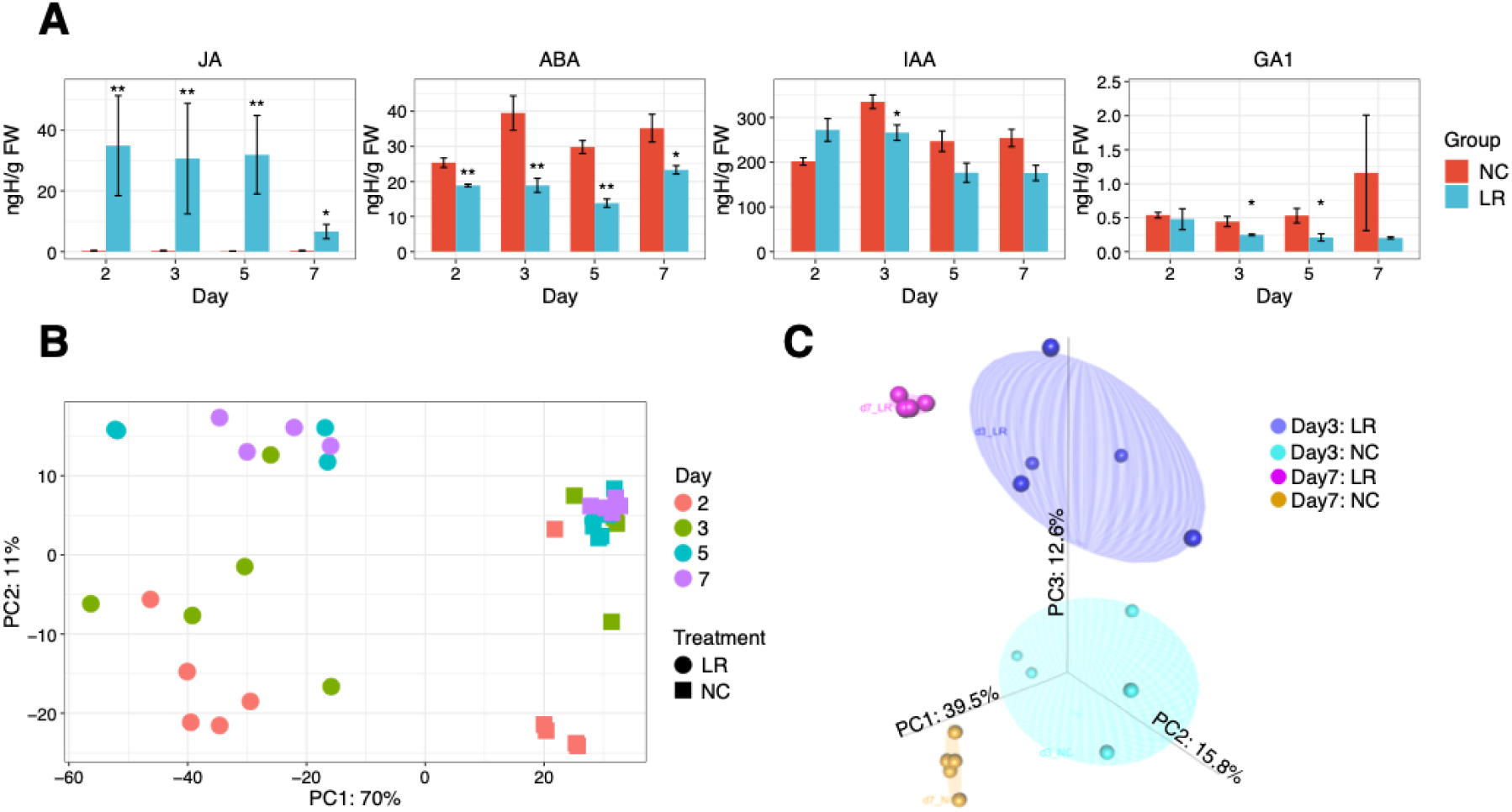
Hormonal and molecular responses of Norway spruce to bark beetle attack. **A** Hormonal changes in Norway spruce in response to bark beetle attack. Mean values and standard errors are represented by bars. Concentrations (ngH/g fresh weight) of four hormones: Jasmonic acid (JA), Abscisic acid (ABA), Indole-3-acetic acid (IAA), and Gibberellin A1 (GA1), in the negative control group (NC) and local response group (LR) at days 2, 3, 5, and 7 after beetle exposure. Significance was determined using Wilcoxon Test (p < 0.05 (*); p < 0.01 (**)). **B** Principal component analysis (PCA) of RNA-seq gene expression data in Norway spruce in response to bark beetle attack. The scatter plot compares the PCA of RNA-seq gene expression profiles, displaying the local response (LR) and negative control group (NC) at days 2, 3, 5, and 7 after exposure. The first principal component (PC1) explains 70% of the variance, while the second principal component (PC2) explains 10%. **C** 3D scatter plot principal component analysis (PCA) of proteomic expression data in Norway spruce following a bark beetle attack. The three-dimensional scatter plot shows the PCA of proteomic profiles, comparing the local response group (LR) and negative control group (NC) at days 3 and 7 after beetle exposure. PC1 explains 39.5% of the variance, PC2 explains 15.8%, and PC3 explains 12.6%. At day 3, a higher degree of variation between the samples was found in contrast to day 7 that shows a more homogeneous protein expression profile.

### RNA-Seq analysis uncovers strong differential gene expression in the local induced response

A principal component analysis (PCA) on the overall differential gene expression profile from RNA-seq analysis (Figure 3B) highlighted significant differences between the LR and NC groups. Combined, PC1 and PC2 captured 81% of the total variance, with PC1 explaining 70% of the variance over the entire observation period. A total of 1,431, 1,912, 1,234, and 1,155 genes were differentially expressed at days 2, 3, 5, and 7, respectively, with an average of 64±4% being over-expressed and 36±4% under-expressed (Supplementary Table 4). Differences between these time points were evident along PC2, accounting for 11% of the variance. A consistent subset of 455 differentially expressed genes (DEGs) were shared across all time points, representing approximately 33±7% of the total DEGs. Enriched Gene Ontology (GO) terms were identified in both over- and under-expressed gene sets across all time points (Supplementary Table 5). In the over-expressed gene sets, several biological processes stood out in the context of Norway spruce defense response against bark beetle attacks, including the chitin catabolic, biosynthetic, isoprenoid biosynthetic, and lignin catabolic processes. In addition, the over-expressed gene set contained more molecular function terms than the under-expressed gene set, with 28 terms identified in the former and 7 in the latter. The over-expressed set was led by ammonia-lyase activity, while the under-expressed set was dominated by hexosyltransferase activity. Many of the terms in the over-expressed gene set were related to plant defense via secondary metabolism, affecting pathways such as phenylpropanoid biosynthesis (e.g., heme binding), flavonoid biosynthesis (e.g., acyltransferase activity), and terpenoid biosynthesis (e.g., magnesium ion binding) (Liu et al., 2019; Yue et al., 2016). With the emphasis on biochemical pathways, Kyoto Encyclopedia of Genes and Genomes (KEGG) annotation revealed several upregulated pathways involved in Norway spruce defense against biotic invaders, with a role in signal transduction and environmental adaptation (plant-pathogen interaction, MAPK signaling, and plant hormone signal transduction), amino acid metabolism (phenylalanine, tyrosine, and tryptophan biosynthesis), and biosynthesis of plant secondary metabolites (phenylpropanoid, flavonoid, stilbenoid, diarylheptanoid, and gingerol, terpenoid backbone) (Hammerbacher et al., 2019; Mithöfer & Boland, 2012) (Supplementary Table 6). Finally, several anti-fungal/insect defense proteins not annotated on KEGG pathways such defensins, chitinase 2-like, proteinase inhibitors, and PR10 were found to be differentially up-regulated among others (Supplementary Table 4).

### Proteomic analysis identified a localized accumulation of enzymes involved in the biosynthesis of plant secondary metabolites

A total of 3,526 proteins, with at least two identified peptides per protein were quantified across all samples collected from both NC and LR groups at days 3 and 7. Day 3 was selected because it exhibited the highest number of DEGs, providing a snapshot of peak molecular activity, while day 7 was chosen to capture potential protein-level changes at later response stages, as protein expression typically lags behind gene regulation. The 3D PCA plot shows a clear separation of the protein expression profiles between the attack (LR) and control groups (PC3: 12.6% variance) at the two selected time points (PC1: 39.5% variance; Figure 3C). In detail, at day 3, our results show a total of 67 proteins with increased abundance in the attack group compared to 3 proteins with decreased abundance (36 and 2 of them were supported by RNA-Seq, respectively). Notably, at day 7, there was a significant increase in differential protein expression, with 164 proteins showing increased abundance and 13 showing decreased abundance in the attack group (51 and 2 of them were supported by RNA-Seq, respectively) (Supplementary Table 7). 52 proteins were consistently more abundant at both time points whereas one showed decreased abundance. Most of the over-expressed proteins were enriched for enzymes in the biosynthesis of secondary metabolites related to plant defense, such as phenylpropanoid, flavonoid, and terpenoid biosynthetic pathways (Figure 4), and amino acid metabolism, including phenylalanine, tyrosine, and tryptophan biosynthesis (Supplementary Table 8 and 9), in agreement with the RNA-seq results. In addition, other interesting proteins with no KEGG annotation, such as dirigent and laccases, were found to be over-expressed, contributing to the oxidation of monolignols derived from the phenylpropanoid pathway, a key step in lignin biosynthesis (Al-Khayri et al., 2023). As a result, lignin polymerization was effective and overall higher within the LR samples from day 3 onwards (Figure 4A).

**Fig. 4.**
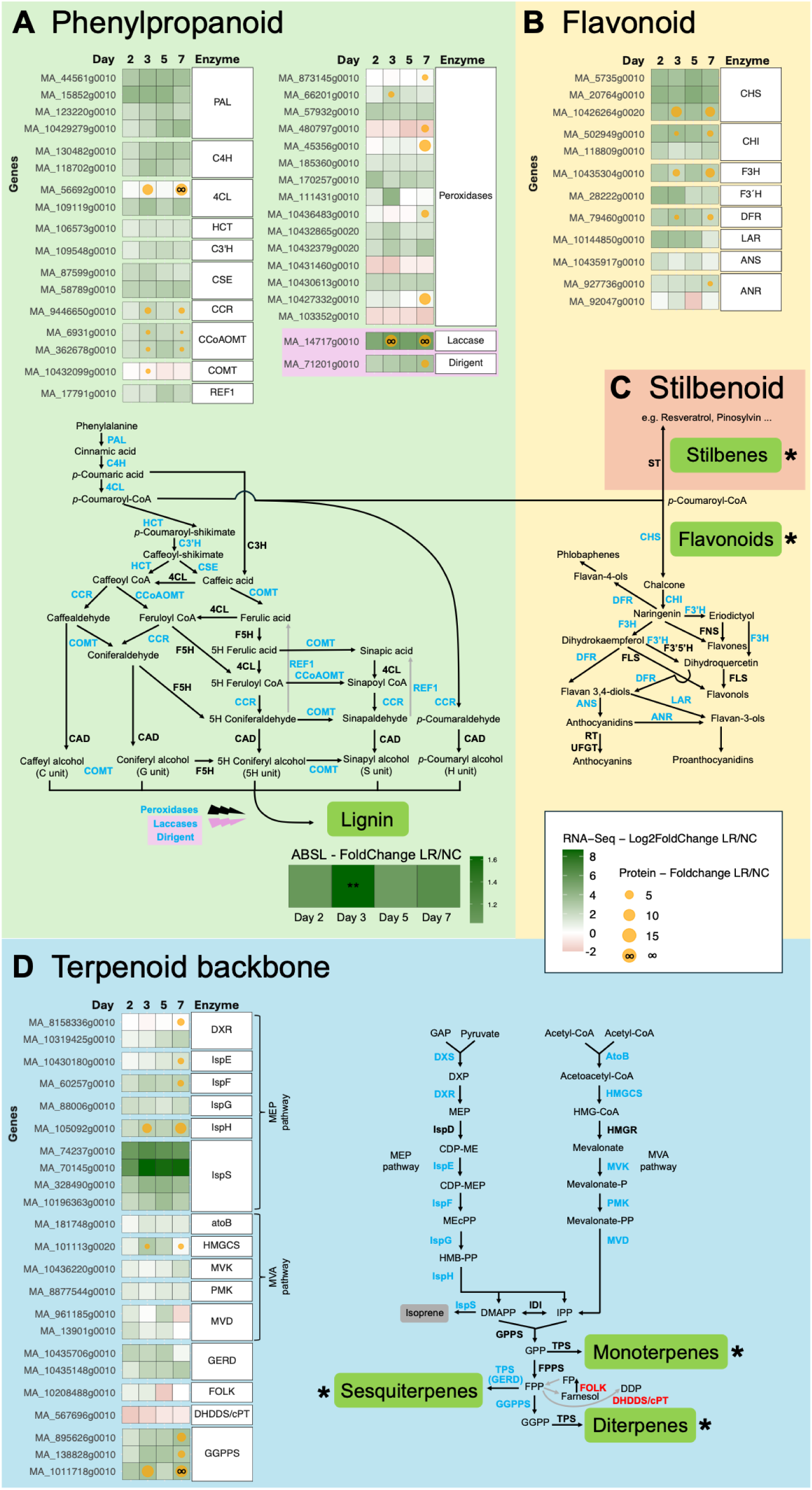
Main genes and key enzymes involved in the KEGG pathways **A** Phenylpropanoid biosynthesis (map00940), **B** Flavonoid biosynthesis (map00941), **C** Stilbenoid biosynthesis (map00945), and **D** Terpenoid backbone biosynthesis (map00900) were differentially upregulated at both the RNA and protein levels (highlighted in blue). This molecular activation was accompanied by increased accumulation of lignin (shown as ABSL), phenolic aglycones, and diterpene resin acids (Figure 5A–C), as well as specific mono- and sesquiterpenoid profiles (Figure 5D–F). An asterisk (*) indicates corresponding metabolomic results presented in Figure 5. For RNA-seq, log₂ fold changes of differentially expressed genes (DEGs) are shown for all timepoints, while proteomic fold changes are presented for days 3 and 7. Only genes and proteins with KEGG annotations and significant differential expression at any timepoint are included. Exceptions were made for Laccase and Dirigent proteins, where representative examples are shown due to the high number of DEGs identified for these crucial enzymes in lignin biosynthesis. A few downregulated genes are highlighted in red. Pathway figures were adapted from Dong & Lin, (2021) and Lin & Pakrasi, (2019).

### Bark beetle attack induces locally elevated levels of phenolic aglycones, and a specific terpenoid profile in Norway spruce

To further investigate the secondary metabolism responses to the bark beetle attack over the course of the trial, targeted LC-MS profiling was performed in all samples from the LR and NC groups, across all collection timepoints. Using an in-house built database, we identified and quantified 33 phenolics and 2 diterpene resin acids. The phenolics compounds belong to the main families of metabolites described in conifers (reviewed in Metsämuuronen & Sirén, 2019), including hydroxybenzoic and hydroxycinnamic acids, flavonoids and stilbenes (Supplementary Table 10). Hierarchical clustering of the 35 metabolites separated them into two clusters (Figure 5A). Interestingly, Cluster 1 metabolites increased in bark beetle-infested trees compared to controls at days 5 and 7, with the combined peak areas showing an increase in total phenolic content (Figure 5B). In contrast to the metabolites in Cluster 2, which were mainly glycosylated (molecules with attached sugar groups), all the compounds in Cluster 1 were aglycons (non-sugar parts derived from glycosides). The increase in aglycons was not mirrored by a decrease in their glycosides, suggesting de novo synthesis of these compounds rather than their release from glycosides by β-glycosidase activity, enhancing the plant’s ability to respond to stressors (Zhang et al., 2021) (Figure 5C).

**Fig. 5.**
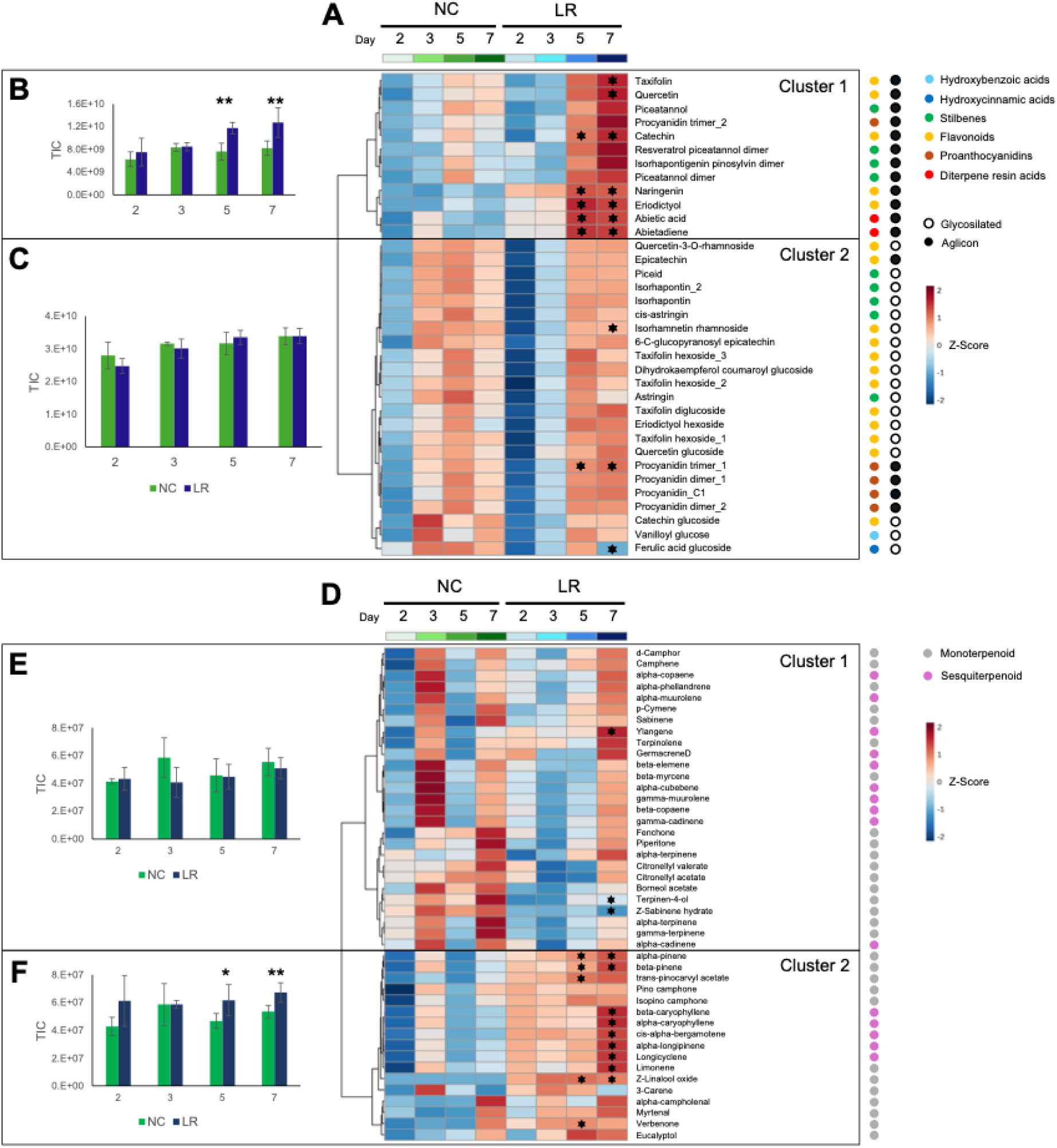
Plant secondary metabolites (PSM) and volatile organic compounds (VOCs) in Norway spruce during bark beetle infestation. **A** Hierarchical clustering of Norway Spruce secondary metabolites during bark beetle infestation. Data were collected from the negative control group (NC) and the local response group (LR) at days 2, 3, 5, and 7 after beetle exposure. Each box represents the average of three biological replicates. Significance was determined using Student’s t-test (black star indicates p < 0.05) between NC and LR samples for the same time point. **B** Total ion current (TIC) for secondary metabolites in Cluster 1. **C** TIC for secondary metabolites in Cluster 2, * indicates p < 0.05 and ** p < 0.01 using Student’s t-test between NC and LR samples for the same time point. **D** Hierarchical clustering of Norway Spruce volatile organic compounds (VOCs) during bark beetle infestation **E** TIC for VOCs in Cluster 1 **F** TIC for VOCs in Cluster 2.

Regarding terpene metabolism, two resin diterpenoids (abietic acid and abietadiene) showed an increase from day 5 onwards (Figure 5A). In addition, a non-targeted VOC profile identified 44 VOCs, 12 hydrocarbon and 17 oxygenated monoterpenoids and 15 sesquiterpenoids (Supplementary Table 11). Hierarchical cluster analyses identified two main clusters with different patterns (Figure 5D). Cluster 1 comprised 27 VOCs that did not change after the attack. Cluster 2 comprised 17 VOCs (e.g., α-pinene, β-pinene, β-caryophyllene, 3-carene, limonene) with a clear tendency towards higher accumulation upon the attack, more evident at day 5 and 7. The combined peak areas of the metabolites from Clusters 1 and 2 showed that terpenoids from Cluster 2 significantly increased upon the attack (Figure 5E and Figure 5F).

## Discussion

In this study we investigated the local molecular defense responses of Norway spruce against pioneer bark beetle attacks using a tailored experimental setup with individual cages for controlled beetle exposure. In an unprecedented multi-omics approach, we provide a comprehensive in situ picture of the tree’s defense mechanisms, revealing coordinated local shifts in phytohormones, and differential gene and protein expression at the sites of attack. These responses were further accompanied by the accumulation of key secondary metabolites, which likely contribute to the formation of a biochemical barrier capable of deterring pioneer beetle colonization. The low attack pressure achieved by this method allowed the induced molecular defense response to be targeted, preventing the depletion of plant resources often observed during natural mass attacks. Such density- and time-dependent studies provide valuable insights to better understand plant-insect interactions and tree resistance to biotic stressors.

Due to its complex genome, *P. abies* serves as a valuable model for exploring plant defenses in conifers (Nystedt et al., 2013). Here, we observed a rapid activation of highly localized defenses (LR) via RT-qPCR (Singh et al., 2024; Trujillo-Moya et al., 2022), suggesting that Norway spruce prioritizes resource allocation to directly attacked tissues during initial beetle boring attempts (Lieutier et al., 2004; Raffa et al., 2005). In contrast, systemic responses (SR) across distal tissues remained unchanged, possibly due to the short 7-day exposure or the limited attack intensity (Netherer et al., 2021). By limiting the early defense responses to the site of first attacks, spruce trees reduce the risk of further infestation while conserving energy for potential future threats. Over evolutionary timescales, such strategies have been employed to balance immediate, highly effective defense needs against long-term survival, fitness, and resource management (Erbilgin et al., 2017). This can be especially advantageous under low beetle densities, where reducing aggregation signals can prevent the escalation of the attack.

A comprehensive gene expression analysis of the local response (LR) using RNA-Seq identified a wide range of genes and pathways involved in the defense process, notably plant-pathogen interaction, MAPK signaling, and plant hormone transduction pathways. Energy metabolism was rapidly impacted after the attack, and the downregulation of photosynthesis-related genes supports the hypothesis of an energy trade-off, shifting resources from growth to defense (Herms & Mattson, 1992). While this may compromise long-term development, it enhances immediate survival and resistance against biotic stressors (Huot et al., 2014). Similar shifts have been reported in studies simulating attacks via MeJA treatment (Mageroy et al., 2020a). In addition, primary metabolism was also affected, with glycolysis being up-regulated to mobilize reserves for the biosynthesis of phenylpropanoids, flavonoids, and terpenoids (Trujillo-Moya et al., 2020).

Differentially expressed genes in the plant-pathogen interaction pathway found in our dataset were predominantly linked to pattern-triggered immunity (PTI) over the effector-triggered immunity (ETI). This localized and rapid defense response is initiated when pathogen-associated molecular patterns (PAMPs) derived from the beetle, are recognized by pattern recognition receptors (PRRs) on the plant cells surface (Nishad et al., 2020). Recognition of PAMPs triggers an influx of calcium ions (Ca²⁺) into the cytoplasm, which is sensed by calmodulin-like proteins (CMLs) and also contributes to the activation of respiratory burst oxidase homologs (RBOHs), both found up-regulated here. RBOHs are enzymes that produce reactive oxygen species (ROS), which act as both signaling molecules and antimicrobial agents by creating a hostile oxidative environment that inhibits pathogen growth (Wu et al., 2023). Meanwhile, CMLs activate a downstream MAPK cascade, contributing to the amplification of immune signaling (Pieterse et al., 2012; Rodriguez et al., 2010). Within this cascade, upregulated MPK6, is a key amplifier of defense signaling and is involved in JA biosynthesis through the observed upregulation of the octadecanoid pathway (Liu & Timko, 2021). JA is subsequently conjugated to isoleucine by the enzyme JAR1, also upregulated here, producing the bioactive form (JA-Ile) that binds to the COI1-JAZ receptor complex (Li et al., 2022). This interaction leads to the degradation of JAZ repressors and the release of MYC2. Here found up-regulated, MYC2 activates the expression of numerous defense-related genes, including those responsible to produce secondary metabolites (terpenoids and phenolics), protease inhibitors, and endochitinases (De Geyter et al., 2012). These defense compounds and proteins might impair the beetle’s ability to inflict further damage while reinforcing the plant’s structural and chemical defenses.

Interestingly, we found that several JAZ proteins encoding genes were upregulated, similar to the findings of Wilkinson et al., 2022) by MeJA treatment. As suggested by the authors this likely represents a feedback mechanism to prevent overactivation of JA-dependent defenses and balance the trade-off between growth and immunity. Such regulation highlights JA’s central role in coordinating hormonal crosstalk and optimizing resource allocation under stress (Kessler & Baldwin, 2002; Mithöfer & Boland, 2008; Wasternack & Hause, 2013). Supporting this, our hormone profiling showed reduced levels of GA1, IAA, and IP, indicating a shift from growth to defense signaling (Gilroy & Breen, 2022). Furthermore, the reduction in abscisic acid (ABA) levels, which typically suppresses JA responses, ensures that JA signaling dominates, optimizing the tree’s capacity to respond to this biotic stress. Finally, while SA levels remained stable, localized overexpression of PR1 suggests some SA-related signaling, although systemic acquired resistance and ETI appeared limited (Anderson et al., 2004; Vlot et al., 2009). These findings highlight the tree’s fine-tuned hormonal crosstalk, which prioritizes JA-driven localized defenses, such as the biosynthesis of secondary metabolites (Liu & Timko, 2021).

Subsequently, in this study, we observed a considerable up-regulation of genes involved in the biosynthesis of phenylpropanoids (11 enzymes) and flavonoids (12 enzymes), with their shared precursor, phenylalanine (13 enzymes), alongside with terpenoids (14 enzymes). These compounds are essential for the tree’s chemical defenses, reinforcing structural integrity and providing antifeedant and antimicrobial properties that discourage herbivore attacks (Faccoli et al., 2005). Our proteome analysis corroborated these transcriptional responses as most of the enzymes in these pathways were either elevated or uniquely detected in the attacked trees. This transcript–protein coordination is particularly meaningful given the multiple regulatory layers between mRNA transcription and protein accumulation, including translational efficiency, post-translational modifications, and protein turnover (Millward et al., 1973). Changes in protein abundance between day 3 and day 7 post-attack suggest a dynamic shift from an acute “defense surge” to the establishment of a longer-term maintenance phase. This adaptive adjustment likely reflects the tree’s need to balance continued resistance with its broader physiological demands (Hammerschmidt, 2009; Heil & Bostock, 2002).

The observed over-expression of the shikimate pathway both at the RNA an proteomic level, has led to an increase in phenylalanine production. Phenylalanine ammonia-lyase (PAL) then channels this phenylalanine into the phenylpropanoid and flavonoid pathways (Nagel et al., 2022), where most transcripts and enzymes were found up-regulated. The phenylpropanoid pathway is directed towards lignin biosynthesis, aligning with a significant lignin increase on day 3. These observations were consistent with stress-induced lignification responses reported in *Picea koraiensis* (Liu et al., 2022). Transient lignification increase strengthens the bark against beetle damage, but is energy-intensive, requiring plants to balance defense and growth. This surge prioritizes structural reinforcement before reallocating resources to other protective strategies (Ma, 2024). Consistent with these findings, our proteomic data further confirmed elevated levels of peroxidases, laccases, and dirigent proteins, which are key enzymes in lignin polymerization and phenolic defense (Ralph et al., 2006).

In addition to lignin biosynthesis, p-coumaroyl-CoA was also diverted into the flavonoid pathway, as indicated by the early upregulation of chalcone synthase and downstream genes and proteins, including FH3 as previously reported Hammerbacher et al. (2019). From day 5 onwards, we observed an increased accumulation of catechin, a flavan-3-ol, along with its precursors naringenin and taxifolin, highlighting their important role in the spruce defense response to biotic stress. Interestingly, the production of stilbenes was also enhanced, probably primed by an excess of p-Coumaroyl-CoA, as no stilbene synthase was found to be up-regulated at neither transcript nor protein level. Notably, all compounds where an increase was observed were aglycones, while glycosylated derivatives remained unchanged, which is consistent with reduced hexosyltransferase activity identified in our enrichment analysis. Polyphenolic parenchyma cells are likely involved in this localized response, as they have been previously shown to accumulate catechin and exhibit elevated PAL activity under bark beetle attacks (Li et al., 2012). Specifically, catechin production during infestation enhances the bark integrity (Zhao et al., 2019), which is vital in preventing beetle penetration by increasing the mechanical strength of the bark. When integrated into cell walls, in synergy with taxifolin, it confers antifeedant properties that halt insects from feeding on plant tissues (Barbehenn & Constabel, 2011; Hammerbacher et al., 2019). Additionally, these compounds provide antioxidant protection, reducing oxidative stress during an attack (Bernatoniene & Kopustinskiene, 2018). Finally, increased amounts of resveratrol and its derivatives, were also detected in our attacked trees. These compounds have been associated with antifeedant activity, as inferred from previous gene expression studies (Faccoli & Schlyter, 2007; Netherer et al., 2021).

Transcriptional changes extend to the terpenoid biosynthesis with the activation of both the mevalonate and MEP/DOXP pathways. Our proteomic data supported this hypothesis by revealing increased levels of enzymes such as geranylgeranyl diphosphate synthase, a key catalyst in the production of diterpenoids like abietadiene and abietic acid, both of which are induced upon attack and contribute to the tree’s resistance by deterring beetle feeding (Hall et al., 2013). Mono- and sesquiterpenes were also found to be elevated in the bark of attacked trees, with higher concentrations of the major components of conifer resin such as the monoterpenes α-pinene, β-pinene, and limonene. The increased monoterpenes levels in the bark have been shown to significantly reduce successful attacks, as indicated by low numbers of visible entry holes (Netherer et al., 2021, 2024; Ott et al., 2021; Zhao et al., 2011; Zhu et al., 2020). Consistently, the isomeric forms of α-pinene and limonene have been found to influence beetle aggregation and colonization (Schiebe et al., 2019), with (-)-α-pinene being a precursor of the *I. typographus* pheromone, while (-)-limonene inhibits the growth of fungi symbiotic to the beetles (Hammerbacher et al., 2019; Schiebe et al., 2019). This dual function of limonene suggests that these pathways can be exploited to enhance the tree’s resistance through genetic or chemical induction methods. Other monoterpenes, like delta-3-carene and pinocamphone, also increased upon attack. Delta-3-carene elicits strong antennal responses and inhibits fungal growth, while pinocamphone may disrupt beetle olfactory communication (Netherer et al., 2021; Schiebe et al., 2019). Notably, several additional monoterpenes were promoted following the attack, with Z-linalool oxide being consistently overproduced. While this compound has not yet been described in spruce defense it is known for its insecticidal and repellent properties making it particularly intriguing (Cheynier et al., 2013; Nyasembe et al., 2014, 2015).

Interestingly, several key defense proteins expected to be over-expressed based on RNA-seq analysis, including multiple basic endochitinases, chitinases, and PR10 proteins, did not exhibit significant changes in protein abundance. Additionally, other defense proteins of interest, including PR1, chitinase 2-like and proteinase inhibitors were not detected in our proteomic dataset. This is likely due to intrinsic technical limitations rather than their biological absence. Plant proteomes are characterized by a high dynamic range, where highly abundant proteins can obscure the detection of low-abundance ones. Furthermore, the sensitivity of current LC-MS/MS systems remains a limiting factor in capturing the full proteome, particularly for small and less ionizable proteins. While our cleanup protocols ensured high sample quality, future studies may benefit from complementary extraction methods or targeted proteomics to expand the possible coverage.

The defense mechanisms of Norway spruce against bark beetle attacks involve a complex interplay of anatomical and biochemical adaptations (Mageroy et al., 2020b), and a balance between constitutive and inducible defenses activated upon trigger stimuli (Franceschi et al., 2005; Ott et al., 2021). In conclusion, our study provides clear evidence that Norway spruce trees have specific induced local defense responses to pioneer bark beetle attacks, which are crucial for their survival. Norway spruce trees, revealed elevated JA levels, extensive gene expression, and protein abundance changes, with the accumulation of key secondary metabolites (Figure 6). The rapid genetic and biochemical changes imply molecular recognition of the beetles triggering an effective cascade of defense responses. Accelerating climate change facilitates multiple beetle generations, creating an urgent need for strategies to bolster Norway spruce resistance as large beetle populations at eruptive mass outbreaks can rapidly deplete this induced local defense response. This molecular dissection provides a comprehensive atlas of Norway spruce molecular defense responses against bark beetles, which could support the development of an efficient exome capture approach for GWAS in this specieś large genome. Here we lay a foundation for resistance-focused breeding (Korolyova, et al., 2022a & 2022b) to strengthen Norway spruce forests against this and other escalating threats from insect herbivores and pathogens, while also offering insights on how to protect other conifer species facing similar ecological threats.

**Fig. 6.**
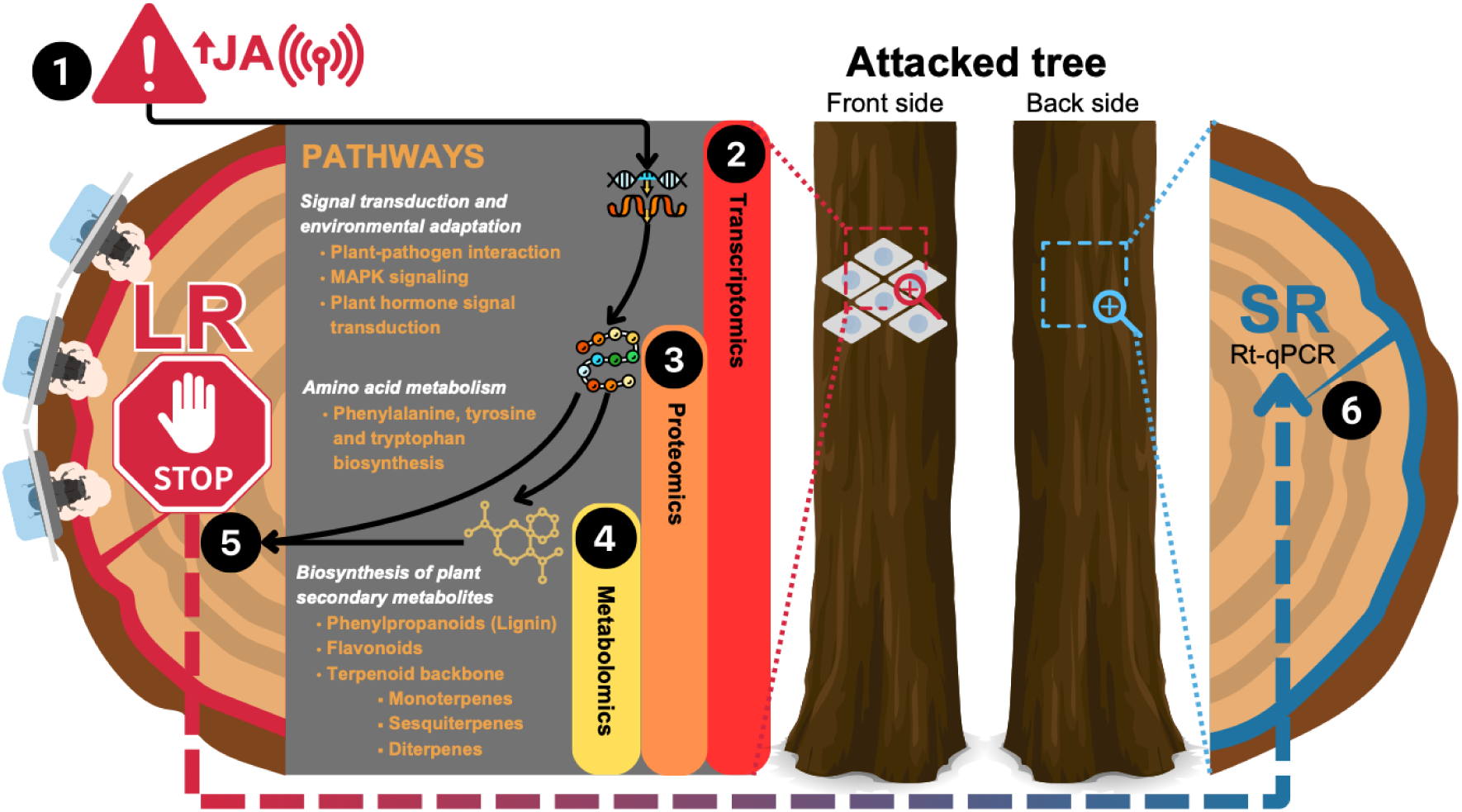
Overview of the local defense response of *Picea abies* to *Ips typographus* attack. The bark beetle is recognized by the tree and JA signaling takes place (1), effectively inducing a differential expression (2) of defense pathways belonging to signal transduction and environmental adaptation, amino acid metabolism, and biosynthesis of plant secondary metabolites. The last two are supported by a proteomic profile (3) that identified key enzymes involved in the production of secondary metabolites (4) such as phenolic aglycones, diterpene resin acids, and specific mono- and sequiterpenes. In addition, lignification takes place to heal the damage, and all defense responses combined can deter and/or halt beetle colonization and establishment (5). In the diametrically opposite side of the attacked area, no systemic response was detected by RT-qPCR (6).

## Materials and methods

### Study design and implementation

The bark beetles used in this study were obtained from a permanent rearing facility (BOKU University, Vienna). They were reared in small logs (60 cm length, 20 cm diameter) under controlled conditions of constant 25 °C with a 16-hour light and 8-hour dark cycle. Freshly emerged young adults had completed their maturation feeding and were collected 1-2 days before the experiments. Only male beetles were used (as they are the gallery-establishing sex); determination of sexes was performed as described in Schlyter & Cederholm, (1981).

This study was conducted in a 35-year-old Norway spruce clonal stand in northern Austria (48.49153460N, 14.62334655E). Forty clonal trees were randomly divided into two groups of 20 individuals each: a negative control and an attack group with controlled exposure to *I. typographus* males (Supplementary Figure 3). To ensure a controlled and consistent exposure, each tree in the attack group was exposed from day 0 to day 7 of the experiment to six beetles, each confined in a specially designed cylindrical cage (25 mm diameter) attached to the bark of the trees (so-called capsular pits) (Supplementary Figure 4). The cages were placed facing south-west at shoulder height (approximately 1.5 m above-ground), ensuring exposure to favorable temperature conditions for beetle activity. Once inside the capsular pits, the beetles were able to move freely while being kept in close contact with the tree, giving them the choice to interact with the tree or not; interactions ranged from gnawing on the bark to boring into bark and phloem for the establishment of a mating chamber.

Disc samples targeting both bark and phloem tissue were collected on days 2, 3, 5, and 7 after the attack using a round hole punch (= cork borer, 25 mm diameter) (Supplementary Figure 5). On each sampling day five trees were randomly selected (replicates) from the initial group of 20 trees in each category (negative control and attack group). In the negative control group (NC), five bark/phloem samples per tree were taken at the same height and orientation as the cages in the attack group were placed. For the attack group, after removal of the cages and beetles, local response (LR) sampling focused on those cage-covered areas showing visible signs of beetle-induced damage, specifically scratches and entry holes created during the attack. Detailed quantification of beetle behavior (e.g., feeding vs. gallery establishment) was beyond the scope of this study and not systematically recorded. Only samples showing visible signs of damage were collected, with a minimum of three samples from each attacked tree. Additionally, systemic response (SR) was assessed by collecting five tissue discs per tree, taken from the side opposite to where the beetles were applied, at the same distance between samples as in the LR group. All samples from a tree under each condition were pooled, treating each tree as a biological replicate (n = 5). Due to initial losses of beetles through death, only four trees were attacked by day 7.

Samples were snap-frozen in liquid nitrogen immediately after collection, transported on dry ice and then stored at -80 °C until further processing using an IKA mill (IKA-Werke, Staufen, Germany) to grind the samples to a fine powder. To prevent any heat-induced molecular degradation, the samples were kept in liquid nitrogen both before and during the grinding process. After grinding, the powdered samples were stored at -80 °C until further molecular analysis.

### RNA isolation

RNA was extracted from 50 mg ground phloem tissue powder according to Protocol A of the Spectrum Plant Total RNA Kit including the On-Column DNase I Digest Set (Sigma-Aldrich, St. Louis, MO, USA) with one modification: a total of five washing steps with wash solution 2 were performed. The RNA Clean & Concentrator-5 Kit (Zymo Research, Irvine, CA, USA) was used to remove PCR inhibitors. RNA concentrations and RNA integrity numbers (RIN) were assessed on a 4200 TapeStation with the RNA ScreenTape assay (Agilent, Santa Clara, CA, USA); only samples showing RIN values ≥ 7 were used for further analysis.

### Quantitative reverse transcriptase polymerase chain reaction (RT-qPCR) analysis

Nineteen Norway spruce genes associated with biotic stress were analyzed via RT-qPCR. Gene selection was done based on previous findings from Trujillo-Moya et al., (2020, 2022). Eight primers from the original studies were kept unchanged, five were modified to increase specificity for the target genes, and nine novel primers were designed (Supplementary Table 1). Additionally, degenerate primers were designed to amplify 10 genes of the dirigent protein family pDIR, ensuring a comprehensive coverage of conserved regions across various sequences. This approach allowed us to assess the overall upregulation of genes responsible for stress response in a more cost-efficient way (Supplementary Figure 2). Primer design was performed based on the *P. abies* transcriptome database (https://treegenesdb.org/org/Picea-abies, accessed on 1st June 2023). Primers were designed using the Eurofins Genomics qPCR Primer & Probe Design software (Eurofins, Ebersberg, Germany, https://eurofinsgenomics.eu/de/ecom/tools/qpcr-assay-design/, accessed on 10 June 2023) using default settings. Additionally, the expression stability of six candidate housekeeping genes (HKGs) was tested. The two most suitable HKGs (elongation factor 1-α and polyubiquitin) based on the RefFinder tool (Xie et al., 2023) were chosen for normalization (Schmidt & Gershenzon, 2008; Trujillo-Moya et al., 2020, 2022; Yakovlev et al., 2006, 2014). RT-qPCR was performed as previously described in (Trujillo-Moya et al., 2020). Data normalization and expression analysis were adjusted for efficiencies using the comparative ddCt method. Gene expression was normalized to housekeeping genes and expressed as fold change using the 2−ΔΔCT formula (Livak & Schmittgen, 2001). In this formula, ΔCT was calculated separately for each time point, where ΔCT = CT (a target gene) − CT (a reference gene), followed by ΔΔCT = ΔCT (negative control) − ΔCT (attacked) and finally 2−ΔΔCT to obtain fold change values. Altogether, our RT-qPCR investigations were compliant with the MIQE guidelines (Bustin et al., 2009). Statistical analyses (unpaired t-test with Welch’s correction) were calculated using GraphPad Prism 10.1.2 (GraphPad Software, San Diego, CA, USA; Supplementary figure 1). PCA plots by using delta Cq values were generated using the R package ggfortify (Tang et al., 2016)

### Identification and quantification of plant hormones

Plant hormone analysis was performed on all LR and NC samples at all four time points. Samples (∼150 mg FW) were suspended in 80% methanol and 1% acetic acid with internal standards and shaken for one hour at 4 °C. The extract was kept at -20 °C overnight and then centrifuged. The supernatant was dried in a vacuum evaporator and redissolved in 1% acetic acid. Purification involved sequential passage through HLB Oasis and MCX columns (Waters), as described in (Seo et al., 2011). The extracts underwent further purification through an ion-exchange WAX column, eluted with 80% methanol-1% acetic acid. The final residues were reconstituted in 5% acetonitrile-1% acetic acid. Hormone separation was performed using UHPLC on a reverse-phase Accucore C18 column, with acetonitrile gradients of 2% to 25% over 13 minutes for cytokinins (CKs) and 2% to 55% over 21 minutes for gibberellins (GAs), indole-3-acetic acid (IAA), abscisic acid (ABA), salicylic acid (SA), and JA. Hormonal analysis was conducted using a Q-Exactive mass spectrometer (Orbitrap detector; Thermo Fisher Scientific, Vienna, Austria) by targeted Selected Ion Monitoring (SIM). The concentrations of hormones in the extracts were determined using embedded calibration curves in Xcalibur 4.0 and TraceFinder 4.1 SP1, using deuterium-labeled standards, except for JA which used dhJA (OlChemim Ltd, Olomouc, Czech Republic).

### RNA-Seq and differential expression analysis

Extracted RNA concentration and quality were assessed with NanoDrop 2000c (Thermo Fisher Scientific, Vienna, Austria) and Fragment Analyzer (Agilent, Vienna, Austria) for LR and NC samples. For those, indexed libraries were prepared from 100 ng of RNA using the QuantSeq 3’ mRNA-Seq Library Prep Kit REV (Lexogen, Vienna, Austria) for Illumina (015UG009V0241) following manufacturing standards and analyzed with Fragment Analyzer (Agilent) using the HS-DNA assay. Libraries were quantified using a Qubit dsDNA HS assay kit (Thermo Fisher Scientific, Vienna, Austria) and sequenced on Illumina NovaSeq6000 with a SR100 read mode (Illumina, San Diego, CA, USA). Sequencing quality control of the raw reads was assessed using FastqQC software and adapter sequences were removed with cutadapt (v1.18; Martin, 2011). Alignment to the reference genome of *P. abies* (Nystedt et al., 2013) and read counting were performed using STAR (v2.6.1a; Dobin & Gingeras, 2015) and featureCounts (v1.6.4; Liao et al., 2014), respectively. RSeQC (Wang et al., 2012, 2016) was used to assess the quality of data after alignment. DEGs were identified with DESeq2 (v1.18.1; Love et al., 2014), using NC samples as controls and adjusting p-values for multiple comparisons with the FDR method. Genes were identified as differentially expressed if they exhibited a minimum 2-fold change and had adjusted p-values below 0.05.

### GO enrichment and KEGG pathways analysis

DEGs identified at each time point were subjected to gene ontology (GO) enrichment analysis. GO annotations were obtained from https://plantgenie.org/ and significance was assessed using the R package topGO (v2.48.0; Alexa & Rahnenfuhrer, 2023) with default parameters (algorithm = weight01 and one-sided Fisher exact test). GO terms were considered significantly enriched if p-values were < 0.01. Visualization of GO terms was performed using ggplot2 (v3.42; Wickham, 2009).

For KEGG pathway analysis, we followed the pipeline described in (Trujillo-Moya et al., 2020, 2022). First, DEGs were annotated to KEGG orthology using bidirectional best hit (BBH) by blasting against a database of 67 plant species (Supplementary Table 12). Then, using custom Python scripts, the genes mapped to each pathway were summarized. This was done separately for up- and down-regulated genes at each time point. Same protocols were applied for differentially expressed proteins.

### Total protein extraction and LCMS analysis

Norway spruce protein extraction was performed following the protocol outlined in (Ramires et al., 2025) on LR and NC samples for day 3 and day 7 only. Total protein concentration was determined using the Pierce 660 nm assay reagent (Thermo Fisher Scientific, Vienna, Austria). Protein concentration was adjusted to 0.5 µg µl^-1^ using resolubilization buffer. Sample preparation for mass spectrometry was performed using a modified version of the S-Trap micro spin column digestion protocol of the manufacturer (ProtiFi LLC, New York, US). For reduction and alkylation, 10 µg of protein, 200 mM tris(2-carboxyethyl)phosphine (TCEP) and 800 mM chloroacetamide (CAA) in 100 mM triethylammonium bicarbonate (TEAB) at an amount of 1:20 (v/v) were filled up with 100 mM TEAB to a total volume of 40 µl, and incubated for 30 minutes at 37 °C in the dark. Then, SDS was added to a final concentration of 2%, and the mixture acidified with phosphoric acid to a final concentration of 1%. After adding S-Trap buffer (90% methanol, 100 mM TEAB, six times the sample volume), proteins were loaded onto the trap column by centrifugation at 1,000 g for 1 minute. The column was washed six times with 150 µl S-Trap buffer by centrifugation at 1,000 g for 1 minute to remove SDS, then dried by centrifugation at 4,000 *g* for 1 minute. Overnight digestion at 37 °C was performed with 1 µg Trypsin/Lys C in 20 µl 50 mM TEAB. Peptides were extracted in three consecutive elution steps using 40 µl 50 mM TEAB, 40 µl 0.2% formic acid (FA), and 40 µl 50% acetonitrile (ACN) by centrifugation at 4,000 *g* for 1 minute, respectively. Peptide eluates were evaporated to dryness in a vacuum centrifuge and resuspended in 50 µl 0.1% trifluoroacetic acid (TFA). Peptide clean-up and desalting was performed using Pierce C18 spin tips (Thermo Fisher Scientific, Vienna, Austria) according to the manufacturer’s instructions and evaporated to dryness in a vacuum centrifuge. Before injection, samples were resuspended in 100 µl 0.1% TFA. The injection volume was 3 µl per sample.

Mass spectrometric data acquisition was performed on a Q Exactive HF Orbitrap mass spectrometer (Thermo Fisher Scientific, Vienna, Austria) connected to a nano-HPLC Ultimate 3000 RSLC system (Dionex, Thermo Fisher Scientific, Vienna, Austria). MS data acquisition as well as qualitative and quantitative data analysis using Proteome Discoverer Software (version 2.4.1.15, Thermo Fisher Scientific, Vienna, Austria) and R version 4.0.5 (R Core Team, 2024), were performed as described in Mayr et al., 2024. Spectra were searched against an in-house *P. abies* database (built using data obtained from https://plantgenie.org, accessed on August 24, 2023) and a database including common contaminants (https://www.thegpm.org/crap/, accessed on June 25, 2019) Proteins with more than two identified tryptic peptides were considered to show significant differences in protein abundance levels when displaying a fold change higher/lower than +/−2-fold and an adjusted *p*-value (according to Benjamini & Hochberg, (1995)) lower than 0.05.

### Lignin quantification

Lignin was quantified using the acetyl bromide soluble lignin (ABSL) assay as previously described (Barnes & Anderson, 2017). Briefly, alcohol insoluble residues were prepared from 19.8 ± 0.3 mg of freeze-dried sample powder by twelve successive washing steps, each including vigorous mixing followed by centrifugation (20,000 *g*, 5 min), with 70% ethanol, 1:1 chloroform:methanol, acetone and 90% DMSO. After each washing with acetone, the pellet was fully dried using a vacuum centrifuge (SPD 111V P2, Thermo Scientific). 4 ± 0.2 mg of the resulting alcohol insoluble residues were incubated with 25% acetyl bromide (v/v in glacial acetic acid) for 1 h at 70 °C, then diluted with glacial acetic acid and mixed with sodium hydroxide and hydroxylamine hydrochloride prior to measuring the absorption at 280 nm. An extinction coefficient value of 21.9 L g^-1^ cm^-1^ was used to estimate the lignin content (Hardell et al.,1980).

### Identification and quantification of phenolic compounds and diterpenes

Frozen powder samples of 50 mg were extracted in 500 µL of 75% acetonitrile with 1 ppm genistein as internal standard. The LC-HRMS analysis was performed on an Orbitrap Exploris 120 mass spectrometer coupled to a Vanquish UHPLC system (Thermo Fisher Scientific, Vienna, Austria) using an Acquity PREMIER BEH C18 UPLC column (1.7 µm, 2.1 x 150 mm; Waters Corp., Mildford, MA, USA). Cromatographic conditions and mass spectrometry specifications were as described in Vazquez-Vilar et al. (2023). Peak identification and quantification were performed using TraceFinder and Compound Discoverer 3.3 software (Thermo Scientific, Waltham, MA, USA). Data analysis and statistics were performed using MetaboAnalyst 6.0 (Pang et al., 2024).

### Identification and quantification of volatile organic compounds (VOCs)

Frozen powder samples of 100 mg were extracted and analyzed using headspace solid phase microextraction (HS-SPME) and GC-MS as described in Trujillo-Moya et al., 2022. Chromatograms were processed using MassHunter software (Agilent Technologies, Vienna, Austria). Compounds were identified by comparing each putative compound with those in the NIST2017 mass spectral library or with our in-house database, which was generated using commercially available compounds. Data analysis and visualization were performed using Simca-P software (Umetrics, Sweden) and MetaboAnalyst 6.0 (Pang et al., 2024).

## Supplementary Figures

**Supplementary Figure 1:**
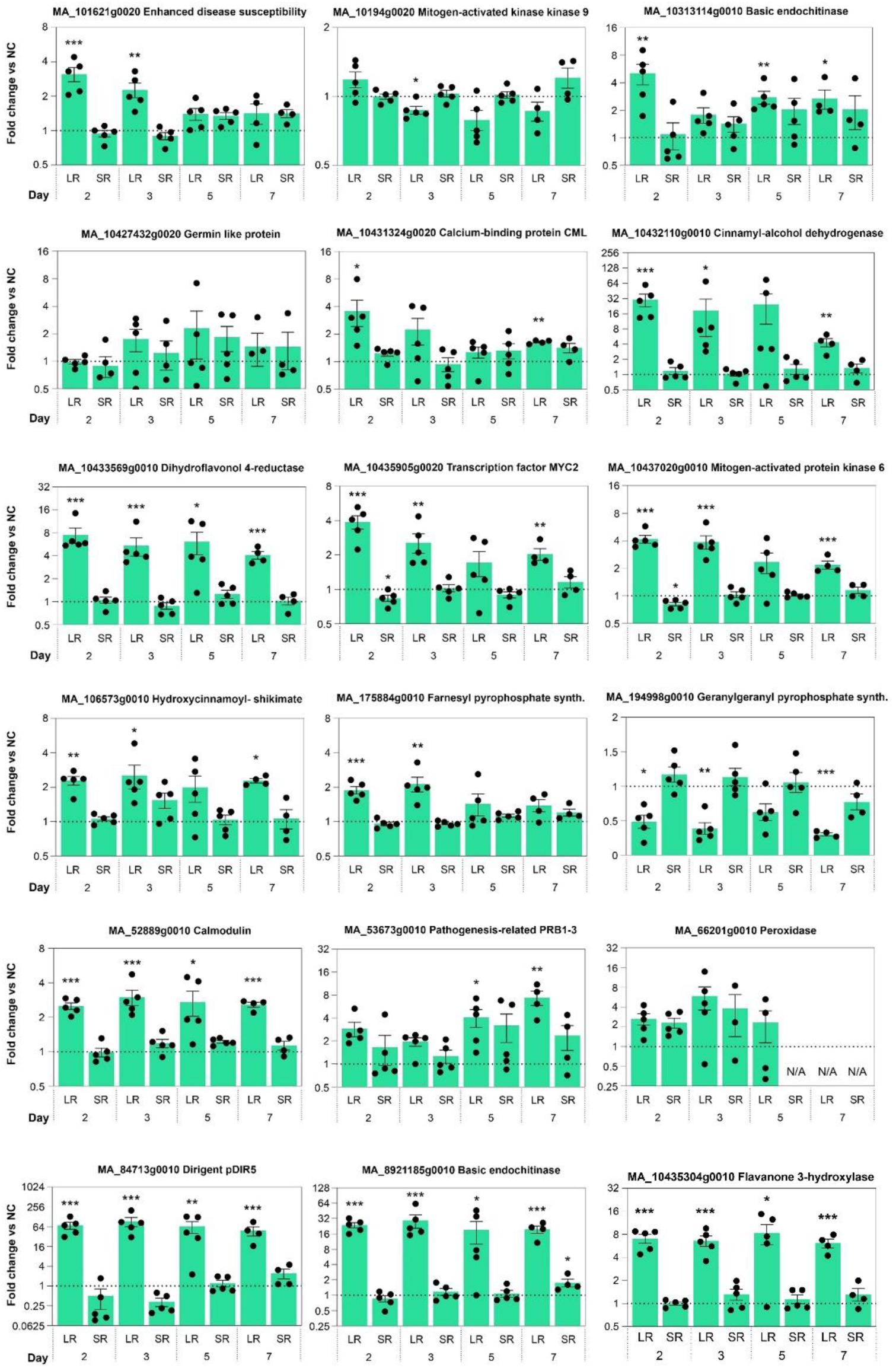
RT-qPCR analysis of defense-related gene expression in *Picea abies*. Statistical significance was determined using an unpaired t-test with Welch’s correction. Significance levels are indicated as follows: p < 0.05 (*), p < 0.01 (**), and p < 0.001 (***).

**Supplementary Figure 2:**
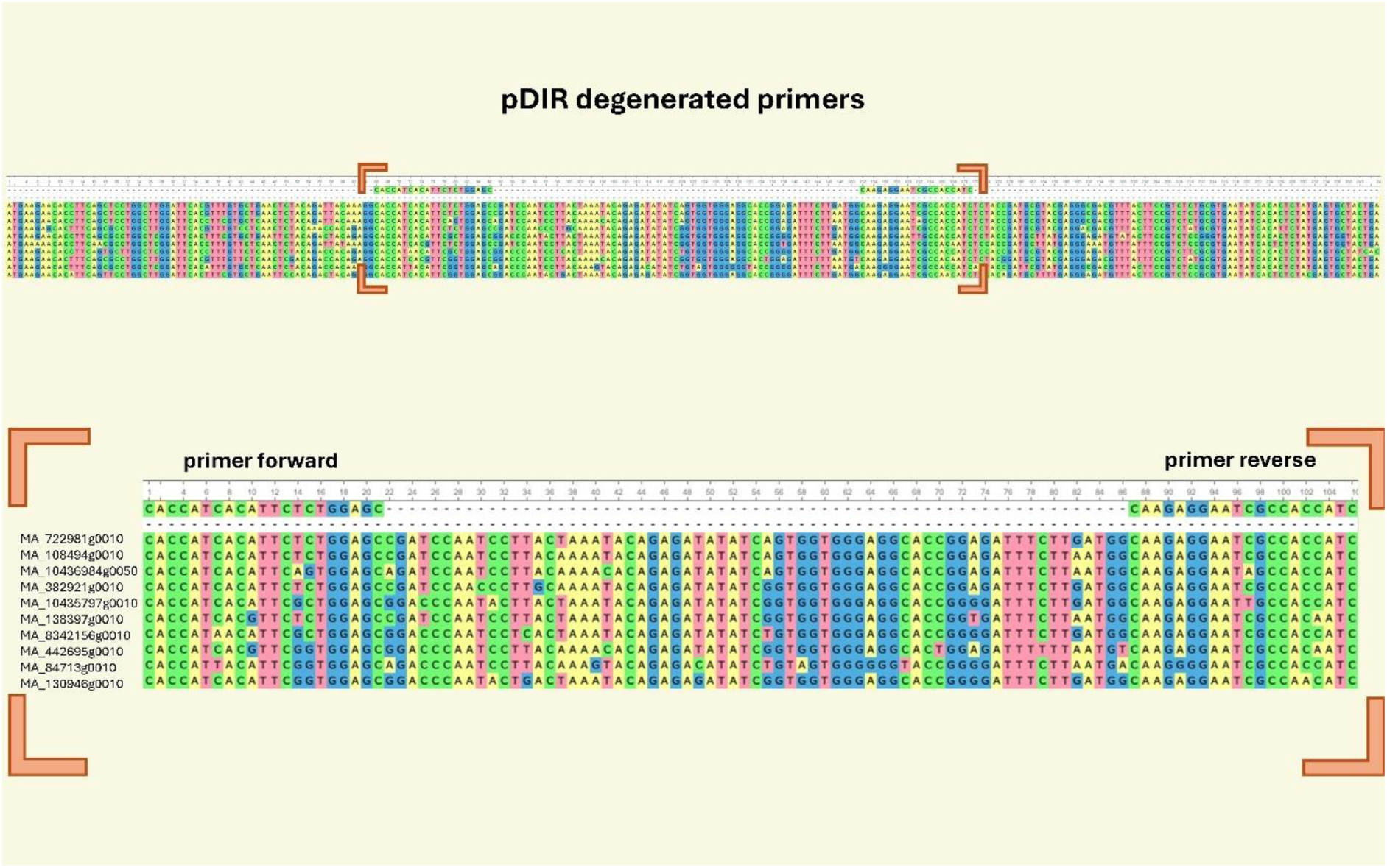
Alignment of target gene sequences and primer binding sites for pDIR degenerated primers.

**Supplementary Figure 3:**
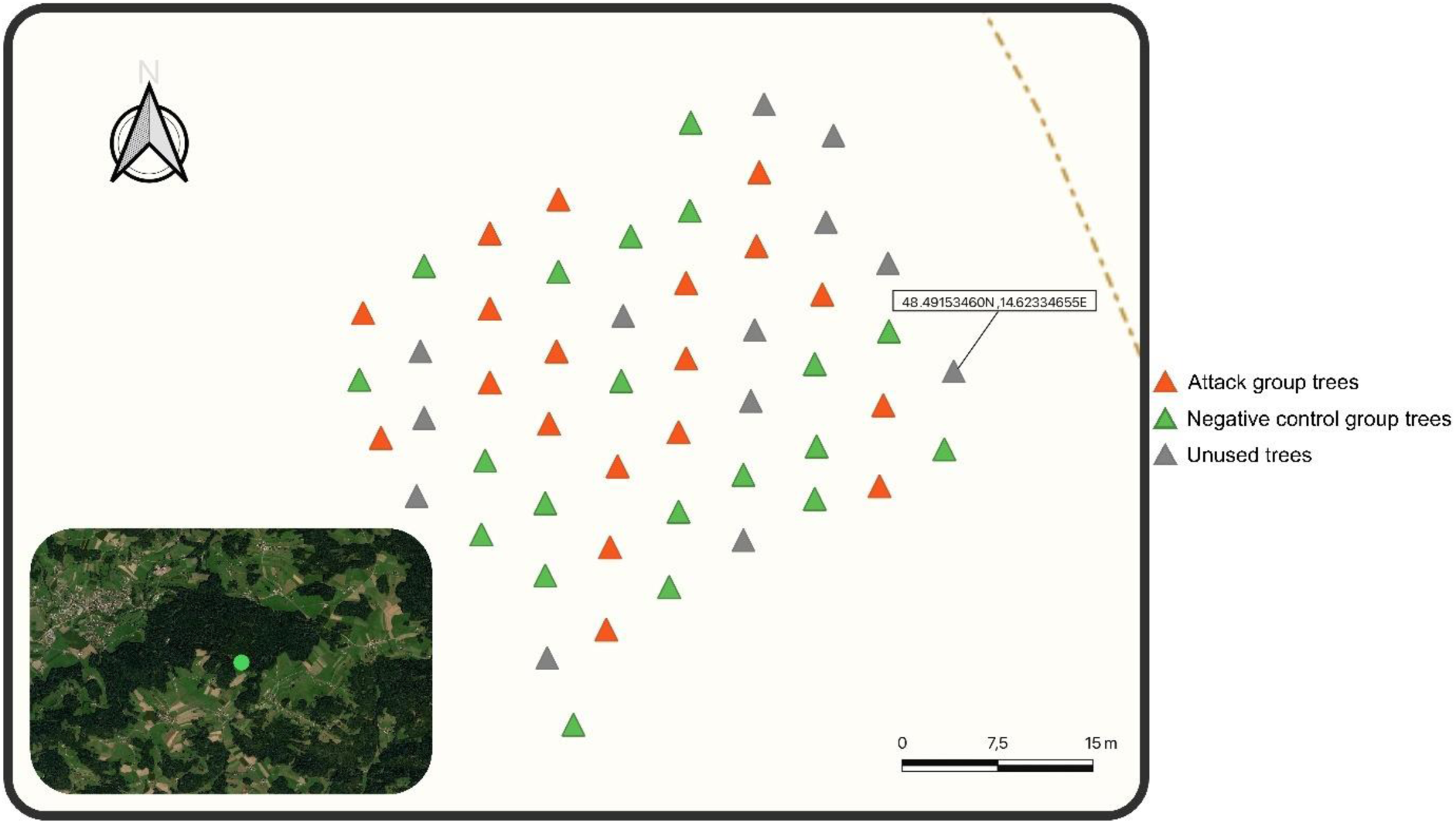
Map of the trial site showing the positions of attack group, negative control group, and unused trees.

**Supplementary Figure 4:**
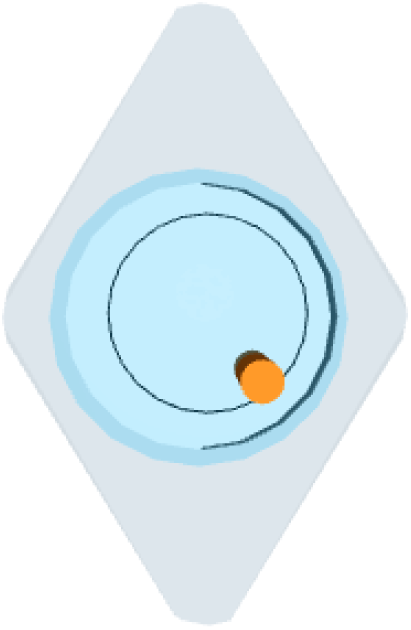
3D render of the capsular pit used to place bark beetles on trees during infestation experiments.

**Supplementary Figure 5:**
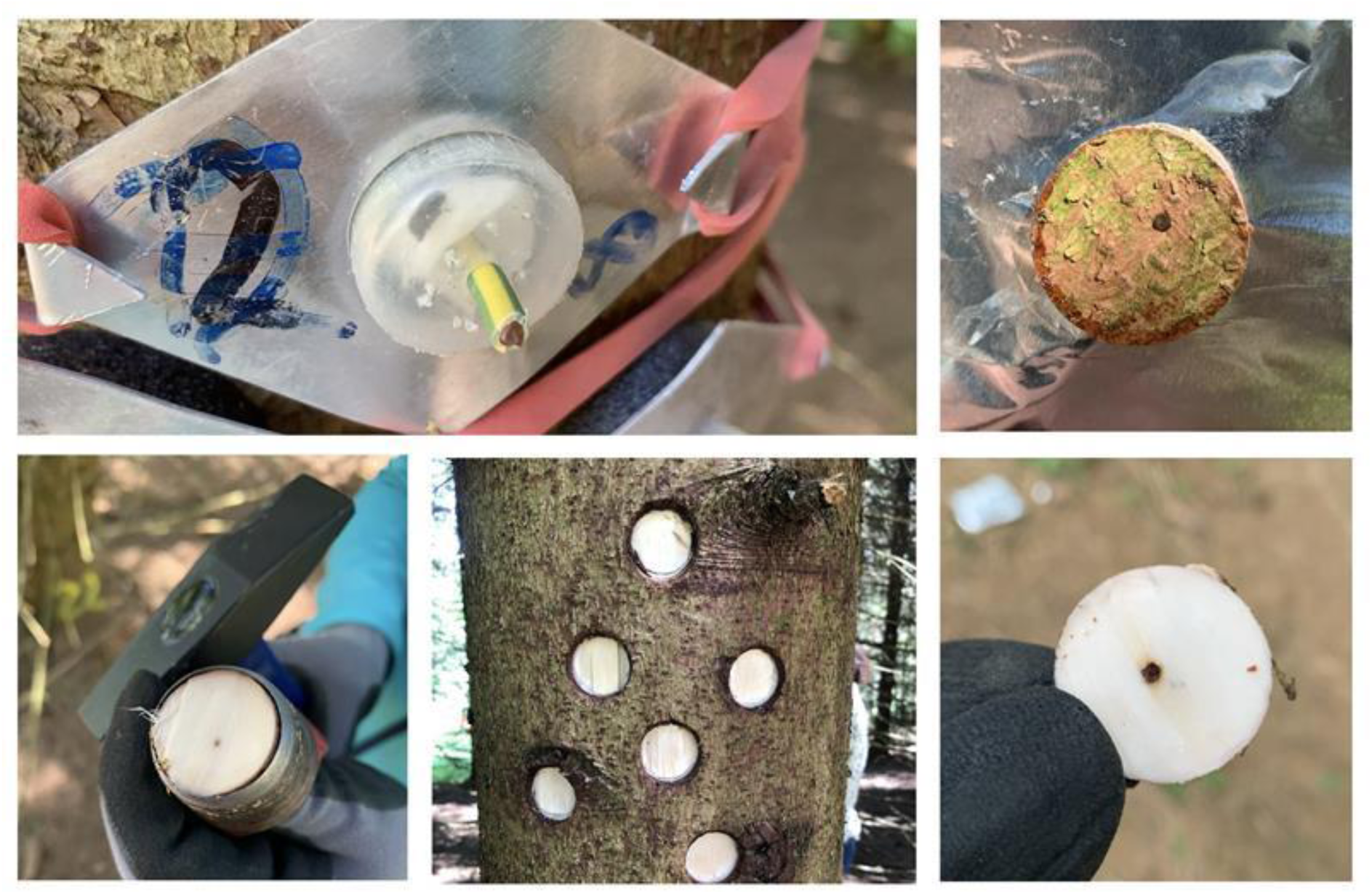
Field setup and sample collection. Bark beetles were confined to the stem using sealed capsular pits. Bark samples were collected from affected tissue for downstream molecular analyses.

## Supplementary Tables

Supplementary Table 1: Primer sequences used for RT-qPCR analysis. The table lists primers used for RT-qPCR in *P. abies*, organized into four categories: Reference genes, Targets – old primers, Targets – remade primers, and Targets – new primers. For each primer pair, the table provides the gene ID, transcript name, forward and reverse sequences (5’–3’), amplicon size (bp), amplification efficiency (E), and the coefficient of determination (R²) from the standard curve.

Supplementary Table 2: Fold change analysis of RT-qPCR data.

Supplementary Table 3: Summary statistics of hormone concentrations (mean ± SE) measured across days and treatments in *Picea abies*. Hormone levels are shown as means with standard errors. Asterisks indicate significant differences between treatments (LR vs. NC) at each time point (Day), based on Wilcoxon rank-sum tests (p < 0.05). Non-significant comparisons are unlabeled.

Supplementary Table 4: DEGs from RNA-Seq and KEGG annotation.

Supplementary Table 5: GO enrichment analysis of DEGs in *P. abies* attacked by *I. typographus*.

Supplementary Table 6: KEGG pathways of DEGs by *P. abies* attacked by *I. typographus*. Numbers refer to key enzymes represented by *P. abies* KAAS assigned orthologous.

Supplementary Table 7: **A** Label-free quantification (LFQ) of proteins across experimental conditions with LCMS/MS. **B** KEGG annotation for differentially expressed proteins.

Supplementary Table 8: GO enrichment analysis of differentially expressed proteins in *P. abies* attacked by *I. typographus*.

Supplementary Table 9: KEGG pathways of differentially expressed proteins by *P. abies* attacked by *I. typographus*. Numbers refer to key enzymes represented by *P. abies* KAAS assigned orthologous.

Supplementary Table 10: Differential accumulation of secondary metabolites across experimental conditions.

Supplementary Table 11: Differential emission of volatile organic compounds (VOCs) under experimental conditions

Supplementary Table 12: Sequence datasets of plant species applied for KEGG Orthology (KO) assignment using the KEGG Automatic Annotation Server (KAAS).

## Data Availability

Raw sequence data have been submitted to the NCBI Short Read Archive (SRA) under BioProject ID: PRJNA1302012

The mass spectrometry proteomics data has been deposited to the ProteomeXchange Consortium via the PRIDE (Perez-Riverol et al., 2022) partner repository with the dataset identifier: PXD066644 and 10.6019/PXD066644.

For data requests, please contact the corresponding author, Dr. Carlos Trujillo-Moya.

## Acknowledgments

The authors thank Otto Dämon, Christine Rossetti and Bettina Lehr for assistance with the lignin quantification. The authors acknowledge Michaella Breuer for her help on the field work. Finally, the authors would like to thank Lila Afifi for her review of the last version and her efforts to improve the grammar and organization of the paper.

## Author contributions

The project was conceptualized by CTM. The research was designed by MR, SN, MS, MvL, and CTM. Access to the experimental site was provided by TL. Bark beetle rearing and cage setup carried out by SN and MS. The experiment was conducted by MR, SN, MS, and MvL. Sample processing was performed by MR. RNA isolation and RT-qPCR analysis were conducted by RE and SS. Plant hormone identification and quantification were performed by EC and JB. GO enrichment analysis, KEGG pathway analysis, and data visualization were carried out by AM. Proteome extraction and LC-MS analysis were performed by MR, KH, SS, and ER-F. Identification and quantification of phenolic compounds, diterpenes, and VOCs were carried out by AE-R, MAM-G and TC. Lignin quantification was performed by EA. Data analysis and initial manuscript drafting were done by MR. Research supervision was provided by CTM and MvL. Project coordination by CTM. All authors edited and approved the manuscript.

## Literature

Alexa, A., & Rahnenfuhrer, J. (2023). Package ‘topGO.’ TopGO: Enrichment Analysis for Gene Ontology. R Package.

Al-Khayri, J. M., Rashmi, R., Toppo, V., Chole, P. B., Banadka, A., Sudheer, W. N., Nagella, P., Shehata, W. F., Al-Mssallem, M. Q., Alessa, F. M., Almaghasla, M. I., & Rezk, A. A. S. (2023). Plant Secondary Metabolites: The Weapons for Biotic Stress Management. In Metabolites (Vol. 13, Issue 6). MDPI. 10.3390/metabo13060716

Allen, C. D., Macalady, A. K., Chenchouni, H., Bachelet, D., McDowell, N., Vennetier, M., Kitzberger, T., Rigling, A., Breshears, D. D., Hogg, E. H. (Ted), Gonzalez, P., Fensham, R., Zhang, Z., Castro, J., Demidova, N., Lim, J. H., Allard, G., Running, S. W., Semerci, A., & Cobb, N. (2010). A global overview of drought and heat-induced tree mortality reveals emerging climate change risks for forests. Forest Ecology and Management, 259(4), 660–684. 10.1016/j.foreco.2009.09.001

Anderson, J. P., Badruzsaufari, E., Schenk, P. M., Manners, J. M., Desmond, O. J., Ehlert, C., Maclean, D. J., Ebert, P. R., & Kazan, K. (2004). Antagonistic interaction between abscisic acid and jasmonate-ethylene signaling pathways modulates defense gene expression and disease resistance in arabidopsis. Plant Cell, 16(12), 3460–3479. 10.1105/tpc.104.025833

Barbehenn, R. V., & Peter Constabel, C. (2011). Tannins in plant-herbivore interactions. In Phytochemistry (Vol. 72, Issue 13, pp. 1551–1565). 10.1016/j.phytochem.2011.01.040

Barnes, W., & Anderson, C. (2017). Acetyl Bromide Soluble Lignin (ABSL) Assay for Total Lignin Quantification from Plant Biomass. BIO-PROTOCOL, 7(5). 10.21769/bioprotoc.2149

Basile, S., Stříbrská, B., Kalyniukova, A., Hradecký, J., Synek, J., Gershenzon, J., & Jirošová, A. (2024). Physiological and biochemical changes of Picea abies (L.) during acute drought stress and their correlation with susceptibility to Ips typographus (L.) and I. duplicatus (Sahlberg). Frontiers in Forests and Global Change, 7. 10.3389/ffgc.2024.1436110

Benjamini, Y., & Hochberg, Y. (1995). Controlling the False Discovery Rate: A Practical and Powerful Approach to Multiple Testing. In Source: Journal of the Royal Statistical Society. Series B (Methodological) (Vol. 57, Issue 1).

Bentz, B. J., Jönsson, A. M., Schroeder, M., Weed, A., Wilcke, R. A. I., & Larsson, K. (2019). Ips typographus and Dendroctonus ponderosae Models Project Thermal Suitability for Intra- and Inter-Continental Establishment in a Changing Climate. Frontiers in Forests and Global Change, 2. 10.3389/ffgc.2019.00001

Bernatoniene, J., & Kopustinskiene, D. M. (2018). The Role of Catechins in Cellular Responses to Oxidative Stress. In Molecules (Vol. 23, Issue 4). MDPI AG. 10.3390/molecules23040965

Bustin, S. A., Benes, V., Garson, J. A., Hellemans, J., Huggett, J., Kubista, M., Mueller, R., Nolan, T., Pfaffl, M. W., Shipley, G. L., Vandesompele, J., & Wittwer, C. T. (2009). The MIQE guidelines: Minimum information for publication of quantitative real-time PCR experiments. Clinical Chemistry, 55(4), 611–622. 10.1373/clinchem.2008.112797

Chakraborty, D., Ciceu, A., Ballian, D., Benito Garzón, M., Bolte, A., Bozic, G., Buchacher, R., Čepl, J., Cremer, E., Ducousso, A., Gaviria, J., George, J. P., Hardtke, A., Ivankovic, M., Klisz, M., Kowalczyk, J., Kremer, A., Lstibůrek, M., Longauer, R., … Schueler, S. (2024). Assisted tree migration can preserve the European forest carbon sink under climate change. Nature Climate Change, 14(8), 845–852. 10.1038/s41558-024-02080-5

Chakraborty, D., Móricz, N., Rasztovits, E., Dobor, L., & Schueler, S. (2021). Provisioning forest and conservation science with high-resolution maps of potential distribution of major European tree species under climate change. Annals of Forest Science, 78(2). 10.1007/s13595-021-01029-4

Cheynier, V., Comte, G., Davies, K. M., Lattanzio, V., & Martens, S. (2013). Plant phenolics: Recent advances on their biosynthesis, genetics, andecophysiology. In Plant Physiology and Biochemistry (Vol. 72, pp. 1–20). 10.1016/j.plaphy.2013.05.009

Clark, E., Huber, D., & Carroll, A. (2012). The Legacy of Attack: Implications of High Phloem Resin Monoterpene Levels in Lodgepole Pines Following Mass Attack by Mountain Pine Beetle, Dendroctonus ponderosae Hopkins. Environmental Entomology, 41, 392–398. 10.1603/EN11295

Cognato, A. I. (2015). Chapter 9 - Biology, Systematics, and Evolution of Ips. In F. E. Vega & R. W. Hofstetter (Eds.), Bark Beetles (pp. 351–370). Academic Press. 10.1016/B978-0-12-417156-5.00009-5

De Geyter, N., Gholami, A., Goormachtig, S., & Goossens, A. (2012). Transcriptional machineries in jasmonate-elicited plant secondary metabolism. Trends in Plant Science, 17(6), 349–359. 10.1016/j.tplants.2012.03.001

Dobin, A., & Gingeras, T. R. (2015). Mapping RNA-seq Reads with STAR. Current Protocols in Bioinformatics, 51(1), 11.14.1-11.14.19. 10.1002/0471250953.BI1114S51

Dong, N. Q., & Lin, H. X. (2021). Contribution of phenylpropanoid metabolism to plant development and plant–environment interactions. In Journal of Integrative Plant Biology (Vol. 63, Issue 1, pp. 180–209). Blackwell Publishing Ltd. 10.1111/jipb.13054

Dyderski, M. K., Paź, S., Frelich, L. E., & Jagodziński, A. M. (2018). How much does climate change threaten European forest tree species distributions? Global Change Biology, 24(3), 1150–1163. 10.1111/gcb.13925

Erbilgin, N., Cale, J. A., Hussain, A., Ishangulyyeva, G., Klutsch, J. G., Najar, A., & Zhao, S. (2017). Weathering the storm: how lodgepole pine trees survive mountain pine beetle outbreaks. Oecologia, 184(2), 469–478. 10.1007/s00442-017-3865-9

Erbilgin, N., Krokene, P., Christiansen, E., Zeneli, G., & Gershenzon, J. (2006). Exogenous application of methyl jasmonate elicits defenses in Norway spruce (Picea abies) and reduces host colonization by the bark beetle Ips typographus. Oecologia, 148(3), 426–436. 10.1007/s00442-006-0394-3

Erbilgin, N., Krokene, P., Kvamme, T., & Christiansen, E. (2007). A host monoterpene influences Ips typographus (Coleoptera: Curculionidae, Scolytinae) responses to its aggregation pheromone. Agricultural and Forest Entomology, 9(2), 135–140. 10.1111/j.1461-9563.2007.00329.x

Faccoli, M., Blaženec, M., & Schlyter, F. (2005). Feeding Response to Host and Nonhost Compounds by Males and Females of the Spruce Bark Beetle Ips typographus in a Tunneling Microassay. Journal of Chemical Ecology, 31(4), 745–759. 10.1007/s10886-005-3542-z

Faccoli, M., & Schlyter, F. (2007). Conifer phenolic resistance markers are bark beetle antifeedant semiochemicals. Agricultural and Forest Entomology, 9(3), 237–245. 10.1111/j.1461-9563.2007.00339.x

Franceschi, V. R., Krokene, P., Christiansen, E., & Krekling, T. (2005). Anatomical and chemical defenses of conifer bark against bark beetles and other pests. In New Phytologist (Vol. 167, Issue 2, pp. 353–376). 10.1111/j.1469-8137.2005.01436.x

Gilroy, E., & Breen, S. (2022). Interplay between phytohormone signalling pathways in plant defence - other than salicylic acid and jasmonic acid. Essays in Biochemistry, 66(5), 657–671. 10.1042/EBC20210089

Hall, D. E., Zerbe, P., Jancsik, S., Quesada, A. L., Dullat, H., Madilao, L. L., Yuen, M., & Bohlmann, J. (2013). Evolution of conifer diterpene synthases: Diterpene resin acid biosynthesis in lodgepole pine and jack pine involves monofunctional and bifunctional diterpene synthases. Plant Physiology, 161(2), 600–616. 10.1104/pp.112.208546

Hammerbacher, A., Kandasamy, D., Ullah, C., Schmidt, A., Wright, L. P., & Gershenzon, J. (2019). Flavanone-3-hydroxylase plays an important role in the biosynthesis of spruce phenolic defenses against bark beetles and their fungal associates. Frontiers in Plant Science, 10. 10.3389/fpls.2019.00208

Hammerbacher, A., Ralph, S. G., Bohlmann, J., Fenning, T. M., Gershenzon, J., & Schmidt, A. (2011). Biosynthesis of the major tetrahydroxystilbenes in spruce, astringin and isorhapontin, proceeds via resveratrol and is enhanced by fungal infection. Plant Physiology, 157(2), 876–890. 10.1104/pp.111.181420

Hammerschmidt, R. (2009). Systemic Acquired Resistance. In Advances in Botanical Research (Vol. 51, Issue C, pp. 173–222). 10.1016/S0065-2296(09)51005-1

Hardell, H.-L.; Leary, G. J.; Stoll, M.; Westermark, U. (1980). Variations in Lignin Structure in Defined Morphological Parts of Spruce. Svensk Papperstidning-Nordisk Cellulosa, 83, 44–49.

Heil, M., & Bostock, R. M. (2002). Induced systemic resistance (ISR) against pathogens in the context of induced plant defences. In Annals of Botany (Vol. 89, Issue 5, pp. 503–512). 10.1093/aob/mcf076

Herms, D. A., & Mattson, W. J. (1992). The quarterly review of biology the dilemma of plants: to grow or defend. In Source: The Quarterly Review of Biology (Vol. 67, Issue 3).

Hlásny, T., Zimová, S., Merganičová, K., Štěpánek, P., Modlinger, R., & Turčáni, M. (2021). Devastating outbreak of bark beetles in the Czech Republic: Drivers, impacts, and management implications. Forest Ecology and Management, 490. 10.1016/j.foreco.2021.119075

Howe, M., Yanchuk, A., Wallin, K. F., & Raffa, K. F. (2024). Quantification of heritable variation in multiple lodgepole pine chemical and physical traits that contribute to defense against mountain pine beetle (Dendroctonus ponderosae). Forest Ecology and Management, 553. 10.1016/j.foreco.2023.121660

Huot, B., Yao, J., Montgomery, B. L., & He, S. Y. (2014). Growth-defense tradeoffs in plants: A balancing act to optimize fitness. In Molecular Plant (Vol. 7, Issue 8, pp. 1267–1287). Oxford University Press. 10.1093/mp/ssu049

Jactel, H., Petit, J., Desprez-Loustau, M. L., Delzon, S., Piou, D., Battisti, A., & Koricheva, J. (2012). Drought effects on damage by forest insects and pathogens: A meta-analysis. Global Change Biology, 18(1), 267–276. 10.1111/j.1365-2486.2011.02512.x

Keeling, C. I., & Bohlmann, J. (2006). Genes, enzymes and chemicals of terpenoid diversity in the constitutive and induced defence of conifers against insects and pathogens. In New Phytologist (Vol. 170, Issue 4, pp. 657–675). 10.1111/j.1469-8137.2006.01716.x

Kessler, A., & Baldwin, I. T. (2002). Plant responses to insect herbivory: The emerging molecular analysis. In Annual Review of Plant Biology (Vol. 53, pp. 299–328). 10.1146/annurev.arplant.53.100301.135207

Korolyova, N., Buechling, A., Ďuračiová, R., Zabihi, K., Turčáni, M., Svoboda, M., Bláha, J., Swarts, K., Poláček, M., Hradecký, J., Červenka, J., Němčák, P., Schlyter, F., & Jakuš, R. (2022a). The Last Trees Standing: Climate modulates tree survival factors during a prolonged bark beetle outbreak in Europe. Agricultural and Forest Meteorology, 322. 10.1016/j.agrformet.2022.109025

Korolyova, N., Buechling, A., Lieutier, F., Yart, A., Cudlín, P., Turčáni, M., & Jakuš, R. (2022b). Primary and secondary host selection by Ips typographus depends on Norway spruce crown characteristics and phenolic-based defenses. Plant Science, 321. 10.1016/j.plantsci.2022.111319

Krokene, P. (2015). Conifer Defense and Resistance to Bark Beetles. In Bark Beetles: Biology and Ecology of Native and Invasive Species (pp. 177–207). Elsevier Inc. 10.1016/B978-0-12-417156-5.00005-8

Krokene, P., Nagy, N. E., & Krekling, T. (2008). Traumatic resin ducts and polyphenolic parenchyma cells in conifers. In Induced Plant Resistance to Herbivory (pp. 147–169). Springer Netherlands. 10.1007/978-1-4020-8182-8_7

Lehmanski, L. M. A., Kandasamy, D., Andersson, M. N., Netherer, S., Alves, E. G., Huang, J., & Hartmann, H. (2023). Addressing a century-old hypothesis – do pioneer beetles of Ips typographus use volatile cues to find suitable host trees? New Phytologist, 238(5), 1762–1770. 10.1111/nph.18865

Li, C., Xu, M., Cai, X., Han, Z., Si, J., & Chen, D. (2022). Jasmonate Signaling Pathway Modulates Plant Defense, Growth, and Their Trade-Offs. In International Journal of Molecular Sciences (Vol. 23, Issue 7). MDPI. 10.3390/ijms23073945

Li, S. H., Nagy, N. E., Hammerbacher, A., Krokene, P., Niu, X. M., Gershenzon, J., & Schneider, B. (2012). Localization of Phenolics in Phloem Parenchyma Cells of Norway Spruce (Picea abies). ChemBioChem, 13(18), 2707–2713. 10.1002/cbic.201200547

Liao, Y., Smyth, G. K., & Shi, W. (2014). featureCounts: an efficient general purpose program for assigning sequence reads to genomic features. Bioinformatics, 30(7), 923–930. 10.1093/BIOINFORMATICS/BTT656

Lieutier F, Day KR, Battisti A, Grégoire J-C, & Evans HF. (2004). Host Resistance to Bark Beetles and Its Variations. In Bark and wood boring insects in living trees in Europe - A Synthesis.

Lin, P. C., & Pakrasi, H. B. (2019). Engineering cyanobacteria for production of terpenoids. In Planta (Vol. 249, Issue 1, pp. 145–154). Springer Verlag. 10.1007/s00425-018-3047-y

Liu, H., & Timko, M. P. (2021). Jasmonic acid signaling and molecular crosstalk with other phytohormones. In International Journal of Molecular Sciences (Vol. 22, Issue 6, pp. 1–24). MDPI AG. 10.3390/ijms22062914

Liu, Y., Liu, L., Yang, S., Zeng, Q., He, Z., & Liu, Y. (2019). Cloning, characterization and expression of the phenylalanine ammonia-lyase gene (PaPAL) from spruce Picea asperata. Forests, 10(8). 10.3390/f10080613

Liu, Y., Zhou, Q., Wang, Z., Wang, H., Zheng, G., Zhao, J., & Lu, Q. (2022). Pathophysiology and transcriptomic analysis of Picea koraiensis inoculated by bark beetle-vectored fungus Ophiostoma bicolor. Frontiers in Plant Science, 13. 10.3389/fpls.2022.944336

Livak, K. J., & Schmittgen, T. D. (2001). Analysis of relative gene expression data using real-time quantitative PCR and the 2-ΔΔCT method. Methods, 25(4), 402–408. 10.1006/meth.2001.1262

Love, M. I., Huber, W., & Anders, S. (2014). Moderated estimation of fold change and dispersion for RNA-seq data with DESeq2. Genome Biology, 15(12), 1–21. 10.1186/S13059-014-0550-8/FIGURES/9

Ma, Q. H. (2024). Lignin Biosynthesis and Its Diversified Roles in Disease Resistance. In Genes (Vol. 15, Issue 3). Multidisciplinary Digital Publishing Institute (MDPI). 10.3390/genes15030295

Mageroy, M. H., Christiansen, E., Långström, B., Borg-Karlson, A. K., Solheim, H., Björklund, N., Zhao, T., Schmidt, A., Fossdal, C. G., & Krokene, P. (2020b). Priming of inducible defenses protects Norway spruce against tree-killing bark beetles. Plant Cell and Environment, 43(2), 420–430. 10.1111/pce.13661

Mageroy, M. H., Wilkinson, S. W., Tengs, T., Cross, H., Almvik, M., Pétriacq, P., Vivian-Smith, A., Zhao, T., Fossdal, C. G., & Krokene, P. (2020a). Molecular underpinnings of methyl jasmonate-induced resistance in Norway spruce. Plant Cell and Environment, 43(8), 1827–1843. 10.1111/pce.13774

Marini, L., Økland, B., Jönsson, A. M., Bentz, B., Carroll, A., Forster, B., Grégoire, J. C., Hurling, R., Nageleisen, L. M., Netherer, S., Ravn, H. P., Weed, A., & Schroeder, M. (2017). Climate drivers of bark beetle outbreak dynamics in Norway spruce forests. Ecography, 40(12), 1426–1435. 10.1111/ecog.02769

Martin, D. M., Fäldt, J., & Bohlmann, J. (2004). Functional characterization of nine Norway spruce TPS genes and evolution of gymnosperm terpene synthases of the TPS-d subfamily. Plant Physiology, 135(4), 1908–1927. 10.1104/pp.104.042028

Martin, M. (2011). Cutadapt removes adapter sequences from high-throughput sequencing reads. EMBnet. Journal, 17(1), 10–12. 10.14806/EJ.17.1.200

Mauri, A., Girardello, M., Strona, G., Beck, P. S. A., Forzieri, G., Caudullo, G., Manca, F., & Cescatti, A. (2022). EU-Trees4F, a dataset on the future distribution of European tree species. Scientific Data, 9(1). 10.1038/s41597-022-01128-5

Mayr, A. L., Hummel, K., Leitsch, D., & Razzazi-Fazeli, E. (2024). A Comparison of Bottom-Up Proteomic Sample Preparation Methods for the Human Parasite Trichomonas vaginalis. ACS Omega, 9(8), 9782–9791. 10.1021/acsomega.3c10040

Metsämuuronen, S., & Sirén, H. (2019). Bioactive phenolic compounds, metabolism and properties: a review on valuable chemical compounds in Scots pine and Norway spruce. In Phytochemistry Reviews (Vol. 18, Issue 3, pp. 623–664). Springer Netherlands. 10.1007/s11101-019-09630-2

Miller, D. R., & Borden, J. H. (2000). Dose-dependent and species-specific responses of pine bark beetles (coleoptera: scolytidae) to monoterpenes in association with pheromones. The Canadian Entomologist, 132(2), 183–195. DOI: 10.4039/Ent132183-2

Millward, D. J., Garlick, P. J., James, W. P. T., Nnanyelugo, D. O., & Ryatt, J. S. (1973). Relationship between Protein Synthesis and RNA Content in Skeletal Muscle. Nature, 204–205.

Mithöfer, A., & Boland, W. (2008). Recognition of herbivory-associated molecular patterns. In Plant Physiology (Vol. 146, Issue 3, pp. 825–831). American Society of Plant Biologists. 10.1104/pp.107.113118

Mithöfer, A., & Boland, W. (2012). Plant defense against herbivores: Chemical aspects. In Annual Review of Plant Biology (Vol. 63, pp. 431–450). 10.1146/annurev-arplant-042110-103854

Nagel, R., Hammerbacher, A., Phillips, M. A., & Phillips, M. A. (2022). Bark Beetle Attack History Does Not Influence the Induction of Terpene and Phenolic Defenses in Mature Norway Spruce (Picea abies) Trees by the Bark Beetle-Associated Fungus Endoconidiophora polonica. Frontiers in Plant Science, 13. 10.3389/fpls.2022.892907

Netherer, S., & Hammerbacher, A. (2021). The Eurasian spruce bark beetle in a warming climate: Phenology, behavior, and biotic interactions. In Bark Beetle Management, Ecology, and Climate Change (pp. 89–131). Elsevier. 10.1016/B978-0-12-822145-7.00011-8

Netherer, S., Kandasamy, D., Jirosová, A., Kalinová, B., Schebeck, M., & Schlyter, F. (2021). Interactions among Norway spruce, the bark beetle Ips typographus and its fungal symbionts in times of drought. In Journal of Pest Science (Vol. 94, Issue 3, pp. 591–614). Springer Science and Business Media Deutschland GmbH. 10.1007/s10340-021-01341-y

Netherer, S., Lehmanski, L., Bachlehner, A., Rosner, S., Savi, T., Schmidt, A., Huang, J., Paiva, M. R., Mateus, E., Hartmann, H., & Gershenzon, J. (2024). Drought increases Norway spruce susceptibility to the Eurasian spruce bark beetle and its associated fungi. New Phytologist, 242(3), 1000–1017. 10.1111/nph.19635

Nishad, R., Ahmed, T., Rahman, V. J., & Kareem, A. (2020). Modulation of Plant Defense System in Response to Microbial Interactions. In Frontiers in Microbiology (Vol. 11). Frontiers Media S.A. 10.3389/fmicb.2020.01298

Nyasembe, V. O., Tchouassi, D. P., Kirwa, H. K., Foster, W. A., Teal, P. E. A., Borgemeister, C., & Torto, B. (2014). Development and assessment of plant-based synthetic odor baits for surveillance and control of malaria vectors. PLoS ONE, 9(2). 10.1371/journal.pone.0089818

Nyasembe, V. O., Tchouassi, D. P., Mbogo, C. M., Sole, C. L., Pirk, C., & Torto, B. (2015). Linalool oxide: Generalist plant based lure for mosquito disease vectors. Parasites and Vectors, 8(1). 10.1186/s13071-015-1184-8

Nystedt, B., Street, N. R., Wetterbom, A., Zuccolo, A., Lin, Y. C., Scofield, D. G., Vezzi, F., Delhomme, N., Giacomello, S., Alexeyenko, A., Vicedomini, R., Sahlin, K., Sherwood, E., Elfstrand, M., Gramzow, L., Holmberg, K., Hällman, J., Keech, O., Klasson, L., … Jansson, S. (2013). The Norway spruce genome sequence and conifer genome evolution. Nature, 497(7451), 579–584. 10.1038/nature12211

Ott, D. S., Davis, T. S., & Mercado, J. E. (2021). Interspecific variation in spruce constitutive and induced defenses in response to a bark beetle-fungal symbiont provides insight into traits associated with resistance. Tree Physiology, 41(7), 1109– 1121. 10.1093/treephys/tpaa170

Pang, Z., Lu, Y., Zhou, G., Hui, F., Xu, L., Viau, C., Spigelman, A. F., MacDonald, P. E., Wishart, D. S., Li, S., & Xia, J. (2024). MetaboAnalyst 6.0: towards a unified platform for metabolomics data processing, analysis and interpretation. Nucleic Acids Research. 10.1093/nar/gkae253

Perez-Riverol, Y., Bai, J., Bandla, C., García-Seisdedos, D., Hewapathirana, S., Kamatchinathan, S., Kundu, D. J., Prakash, A., Frericks-Zipper, A., Eisenacher, M., Walzer, M., Wang, S., Brazma, A., & Vizcaíno, J. A. (2022). The PRIDE database resources in 2022: A hub for mass spectrometry-based proteomics evidences. Nucleic Acids Research, 50(D1), D543–D552. 10.1093/nar/gkab1038

Pieterse, C. M. J., Van Der Does, D., Zamioudis, C., Leon-Reyes, A., & Van Wees, S. C. M. (2012). Hormonal modulation of plant immunity. Annual Review of Cell and Developmental Biology, 28, 489–521. 10.1146/annurev-cellbio-092910-154055

R Core Team (2024). _R: A Language and Environment for Statistical Computing_. R Foundation for Statistical Computing, Vienna, Austria.

Raffa, Kenneth F.; Aukema, Briah H.; Erbilgin, Nadir; Klepzig, Kier D.; Wallin, Kimberly F. 2005. Interactions among conifer terpenoids and bark beetles across multiple levels of scale: An attempt to understand links between population patterns and physiological processes. Recent Advances in Phytochemistry 39: 79–118

Raffa, K. F., & Berryman, A. A. (1982). Physiological Differences Between Lodgepole Pines Resistant and Susceptible to the Mountain Pine Beetle 1 and Associated Microorganisms 2. Environmental Entomology, 11(2), 486–492. 10.1093/ee/11.2.486

Raffa, K. F., Grégoire, J. C., & Lindgren, B. S. (2015). Natural History and Ecology of Bark Beetles. In Bark Beetles: Biology and Ecology of Native and Invasive Species (pp. 1–40). Elsevier Inc. 10.1016/B978-0-12-417156-5.00001-0

Ralph, S., Park, J. Y., Bohlmann, J., & Mansfield, S. D. (2006). Dirigent proteins in conifer defense: Gene discovery, phylogeny, and differential wound- and insect-induced expression of a family of DIR and DIR-like genes in spruce (Picea spp.). Plant Molecular Biology, 60(1), 21–40. 10.1007/s11103-005-2226-y

Ramires, M., Schlosser, S., Hummel, K., Razzazi-Fazeli, E., van Loo, M., & Trujillo-Moya, C. (2025). Protocol for Proteome Extraction from Norway Spruce Bark Samples. 10.17504/protocols.io.ewov11y7pvr2/v1

Rodriguez, C. M., Petersen, M., & Mundy, J. (2010). Mitogen-activated protein kinase signaling in plants. Annual Review of Plant Biology, 61, 621–649. 10.1146/annurev-arplant-042809-112252

Rouault, G., Candau, J. N., Lieutier, F., Nageleisen, L. M., Martin, J. C., & Warzée, N. (2006). Effects of drought and heat on forest insect populations in relation to the 2003 drought in Western Europe. In Annals of Forest Science (Vol. 63, Issue 6, pp. 613–624). 10.1051/forest:2006044

Schiebe, C., Unelius, C. R., Ganji, S., Binyameen, M., Birgersson, G., & Schlyter, F. (2019). Styrene, (+)-trans-(1R,4S,5S)-4-Thujanol and Oxygenated Monoterpenes Related to Host Stress Elicit Strong Electrophysiological Responses in the Bark Beetle Ips typographus. Journal of Chemical Ecology, 45(5–6), 474–489. 10.1007/s10886-019-01070-8

Schlyter, F., & Cederholm, I. (1981). Separation of the sexes of living spruce bark beetles, Ips typographus (L.), (Coleoptera: Scolytidae). Zeitschrift Für Angewandte Entomologie, 92(1–5), 42–47. 10.1111/j.1439-0418.1981.tb01650.x

Schmidt, A., & Gershenzon, J. (2008). Cloning and characterization of two different types of geranyl diphosphate synthases from Norway spruce (Picea abies). Phytochemistry, 69(1), 49–57. 10.1016/j.phytochem.2007.06.022

Seidl, R., Müller, J., Hothorn, T., Bässler, C., Heurich, M., & Kautz, M. (2015). Small beetle, large-scale drivers: how regional and landscape factors affect outbreaks of the European spruce bark beetle Europe PMC Funders Group. J Appl Ecol, 53(2), 530–540. 10.5061/dryad.c5g9s

Seo, M., Jikumaru, Y., & Kamiya, Y. (2011). Profiling of Hormones and Related Metabolites in Seed Dormancy and Germination Studies. In A. R. Kermode (Ed.), Seed Dormancy: Methods and Protocols (pp. 99–111). Humana Press. 10.1007/978-1-61779-231-1_7

Singh, V. V., Naseer, A., Sellamuthu, G., Mogilicherla, K., Gebauer, R., Roy, A., & Jakuš, R. (2024). Robust reference gene selection in Norway spruce: essential for real-time quantitative PCR across different tissue, stress and developmental conditions. Frontiers in Forests and Global Change, 7. 10.3389/ffgc.2024.1458554

Smith, T., Rizzo, D. M., & North, M. (2005). Patterns of Mortality in an Old-Growth Mixed-Conifer Forest of the Southern Sierra Nevada.

Tang, Y., Horikoshi, M., & Li, W. (2016). Ggfortify: Unified interface to visualize statistical results of popular r packages. R Journal, 8(2). 10.32614/rj-2016-060

The State of the World’s Forests 2020. (2020). In *The State of the World’s Forests* 2020. FAO and UNEP. 10.4060/ca8642en

Treutter, D. (2006). Significance of flavonoids in plant resistance: A review. In Environmental Chemistry Letters (Vol. 4, Issue 3, pp. 147–157). 10.1007/s10311-006-0068-8

Trujillo-Moya, C., Ganthaler, A., Stöggl, W., Arc, E., Kranner, I., Schueler, S., Ertl, R., Espinosa-Ruiz, A., Martínez-Godoy, M. Á., George, J. P., & Mayr, S. (2022). Advances in understanding Norway spruce natural resistance to needle bladder rust infection: transcriptional and secondary metabolites profiling. BMC Genomics, 23(1). 10.1186/s12864-022-08661-y

Trujillo-Moya, C., Ganthaler, A., Stöggl, W., Kranner, I., Schüler, S., Ertl, R., Schlosser, S., George, J. P., & Mayr, S. (2020). RNA-Seq and secondary metabolite analyses reveal a putative defence-transcriptome in Norway spruce (Picea abies) against needle bladder rust (Chrysomyxa rhododendri) infection. BMC Genomics, 21(1). 10.1186/s12864-020-6587-z

Vazquez-Vilar, M., Fernandez-del-Carmen, A., Garcia-Carpintero, V., Drapal, M., Presa, S., Ricci, D., Diretto, G., Rambla, J. L., Fernandez-Muñoz, R., Espinosa-Ruiz, A., Fraser, P. D., Martin, C., Granell, A., & Orzaez, D. (2023). Dually biofortified cisgenic tomatoes with increased flavonoids and branched-chain amino acids content. Plant Biotechnology Journal, 21(12), 2683–2697. 10.1111/pbi.14163

Vlot, A. C., Dempsey, D. A., & Klessig, D. F. (2009). Salicylic acid, a multifaceted hormone to combat disease. Annual Review of Phytopathology, 47, 177–206. 10.1146/annurev.phyto.050908.135202

Wang, L., Nie, J., Sicotte, H., Li, Y., Eckel-Passow, J. E., Dasari, S., Vedell, P. T., Barman, P., Wang, L., Weinshiboum, R., Jen, J., Huang, H., Kohli, M., & Kocher, J. P. A. (2016). Measure transcript integrity using RNA-seq data. BMC Bioinformatics, 17(1). 10.1186/s12859-016-0922-z

Wang, L., Wang, S., & Li, W. (2012). RSeQC: Quality control of RNA-seq experiments. Bioinformatics, 28(16), 2184–2185. 10.1093/bioinformatics/bts356

Wasternack, C., & Hause, B. (2013). Jasmonates: Biosynthesis, perception, signal transduction and action in plant stress response, growth and development. An update to the 2007 review in Annals of Botany. In Annals of Botany (Vol. 111, Issue 6, pp. 1021–1058). 10.1093/aob/mct067

Wickham, H. (2009). ggplot2. Ggplot2. 10.1007/978-0-387-98141-3

Wilkinson, S. W., Dalen, L. S., Skrautvol, T. O., Ton, J., Krokene, P., & Mageroy, M. H. (2022). Transcriptomic changes during the establishment of long-term methyl jasmonate-induced resistance in Norway spruce. Plant Cell and Environment, 45(6), 1891–1913. 10.1111/pce.14320

Wu, B., Qi, F., & Liang, Y. (2023). Fuels for ROS signaling in plant immunity. Trends in Plant Science, 28(10), 1124–1131. 10.1016/j.tplants.2023.04.007

Xie, F., Wang, J., & Zhang, B. (2023). RefFinder: a web-based tool for comprehensively analyzing and identifying reference genes. Functional & Integrative Genomics, 23. 10.1007/s10142-023-01055-7

Yakovlev, I. A., Fossdal, C. G., Johnsen, Ø., Junttila, O., & Skrøppa, T. (2006). Analysis of gene expression during bud burst initiation in Norway spruce via ESTs from subtracted cDNA libraries. Tree Genetics and Genomes, 2(1), 39–52. 10.1007/s11295-005-0031-z

Yakovlev, I. A., Lee, Y. K., Rotter, B., Olsen, J. E., Skrøppa, T., Johnsen, Ø., & Fossdal, C. G. (2014). Temperature-dependent differential transcriptomes during formation of an epigenetic memory in Norway spruce embryogenesis. Tree Genetics and Genomes, 10(2), 355–366. 10.1007/s11295-013-0691-z

Yue, X., Li, X. G., Gao, X. Q., Zhao, X. Y., Dong, Y. X., & Zhou, C. (2016). The Arabidopsis phytohormone crosstalk network involves a consecutive metabolic route and circular control units of transcription factors that regulate enzyme-encoding genes. BMC Systems Biology, 10(1). 10.1186/s12918-016-0333-9

Zhang, W., Sheng, W., Yang, J., Chuanzhi, K., Huang, L., & Guo, L. (2021). Glycosylation of Plant Secondary Metabolites: Regulating from Chaos to Harmony. Environmental and Experimental Botany, 194, 104703. 10.1016/j.envexpbot.2021.104703

Zhao, T., Kandasamy, D., Krokene, P., Chen, J., Gershenzon, J., & Hammerbacher, A. (2019). Fungal associates of the tree-killing bark beetle, Ips typographus, vary in virulence, ability to degrade conifer phenolics and influence bark beetle tunneling behavior. Fungal Ecology, 38, 71–79. 10.1016/j.funeco.2018.06.003

Zhao, T., Krokene, P., Björklund, N., Långström, B., Solheim, H., Christiansen, E., & Borg-Karlson, A.-K. (2010). The influence of Ceratocystis polonica inoculation and methyl jasmonate application on terpene chemistry of Norway spruce, Picea abies. Phytochemistry, 71(11), 1332–1341. 10.1016/j.phytochem.2010.05.017

Zhao, T., Krokene, P., Hu, J., Christiansen, E., Björklund, N., Långström, B., Solheim, H., & Borg-Karlson, A. K. (2011). Induced terpene accumulation in Norway spruce inhibits bark beetle colonization in a dose-dependent manner. PLoS ONE, 6(10). 10.1371/journal.pone.0026649

Zhu, X., Xu, B., Kader, A., Song, B., Zhang, Z., Li, F., & Yang, S. (2020). Behavioral Responses of Scolytus schevyrewi (Coleoptera: Curculionidae: Scolytinae) to Volatiles from Apricot Tree (Rosales: Rosaceae). Environmental Entomology, 49(3), 586–592. 10.1093/ee/nvaa027

