## Supplementary figures and images for "Pioneer bark beetle attacks induce multifaceted localized defense responses in Norway spruce"

### Supplementary Figure 1

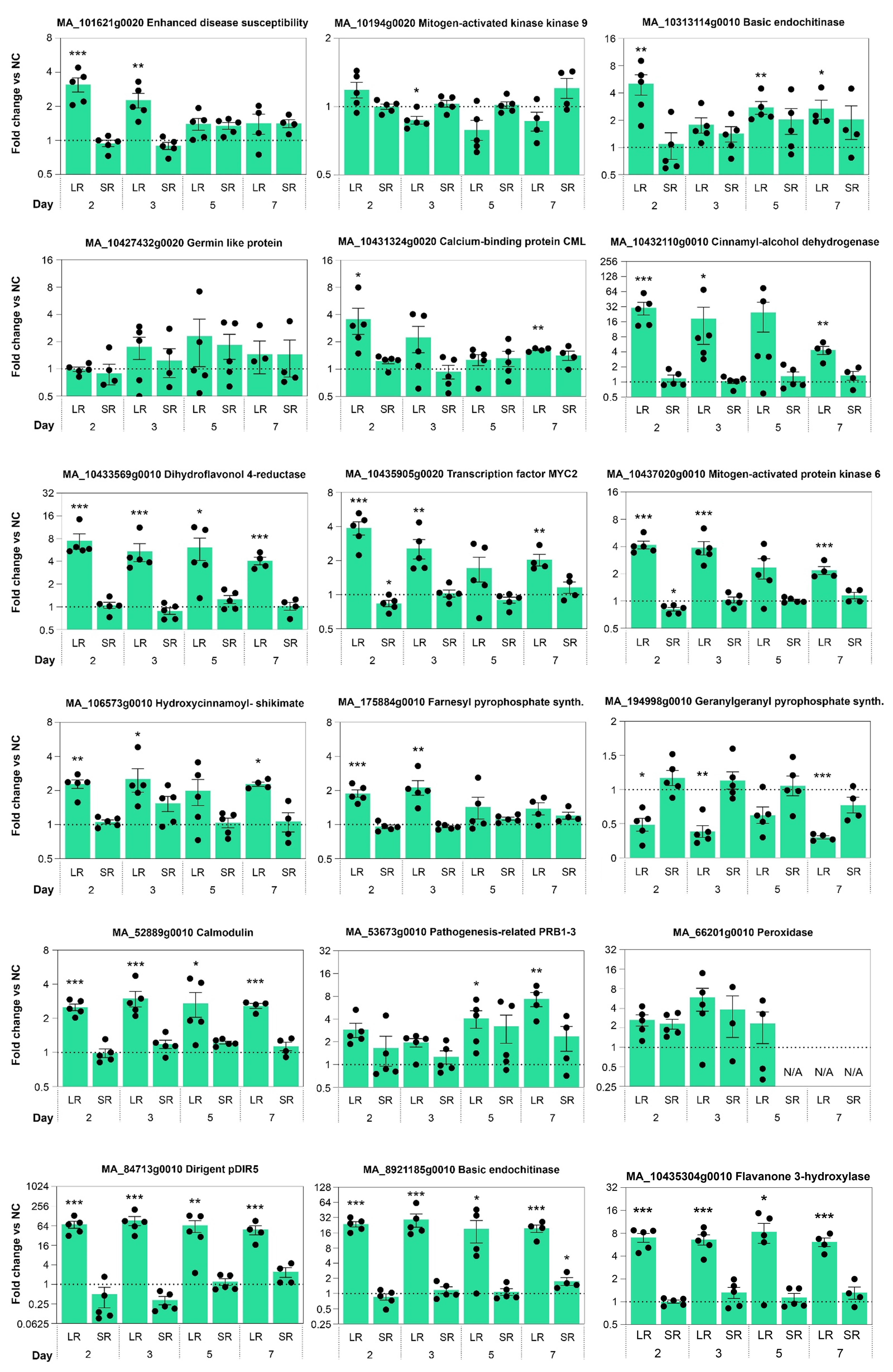

### Supplementary Figure 2

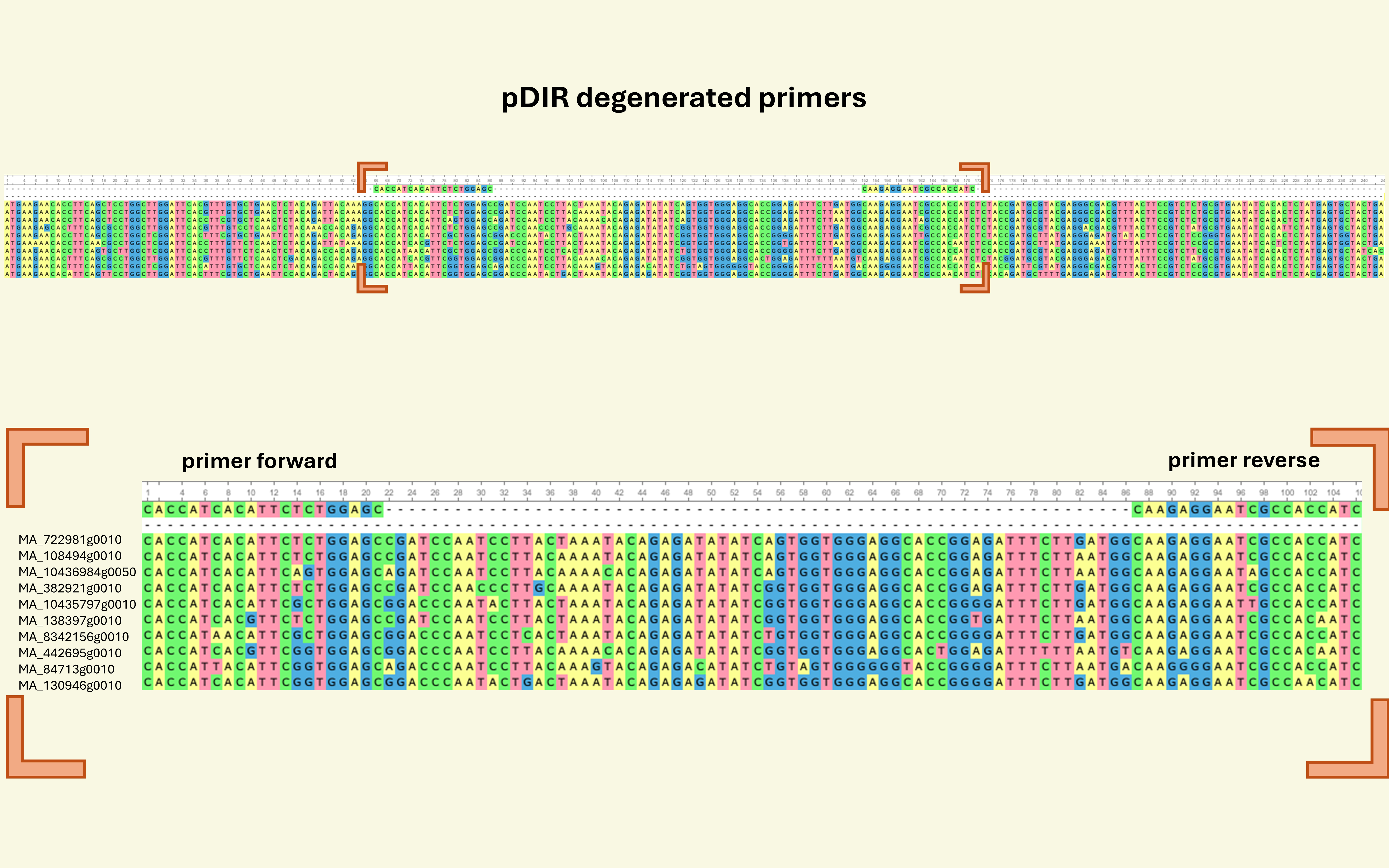

### Supplementary Figure 3

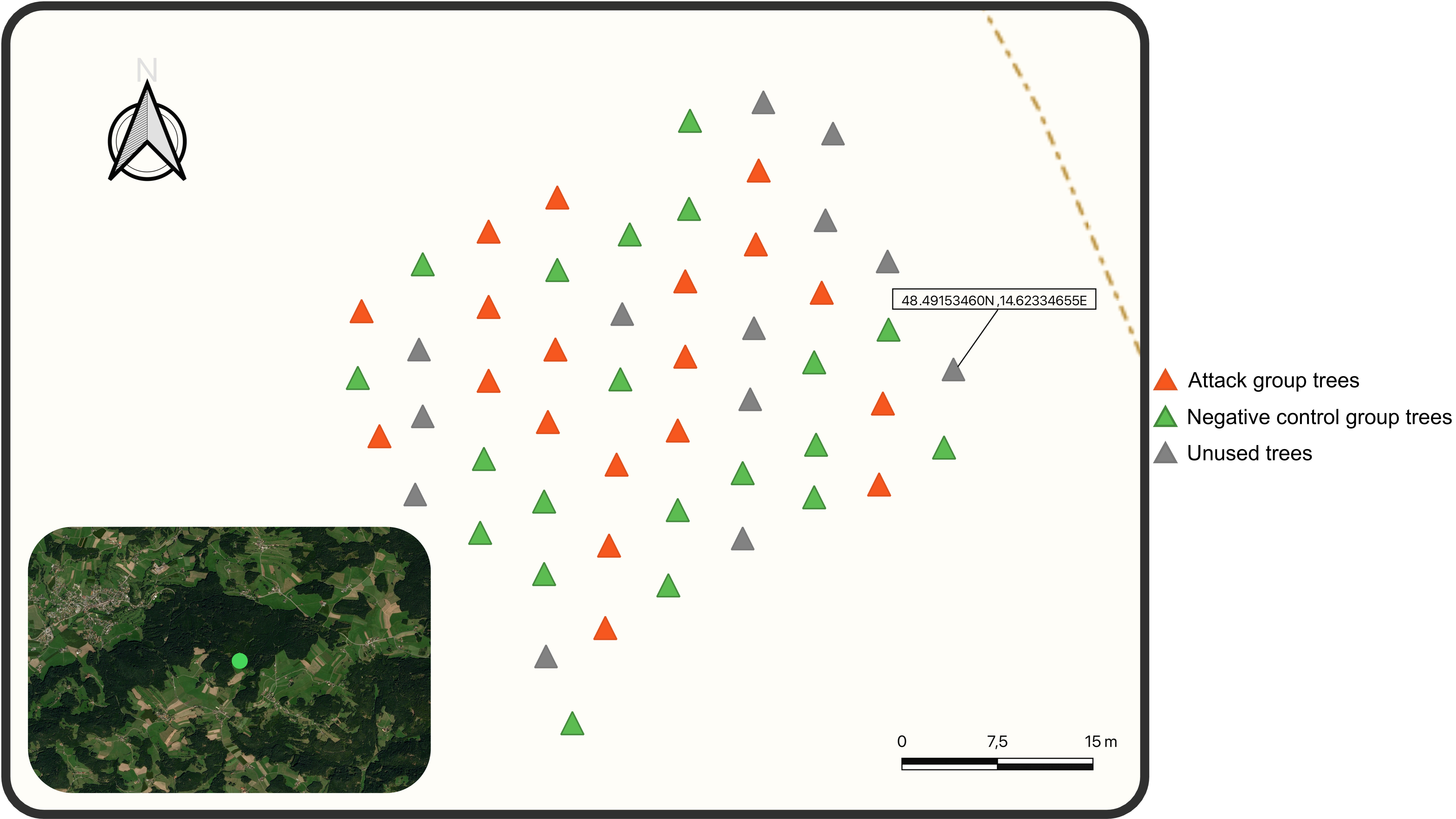

### Supplementary Figure 4

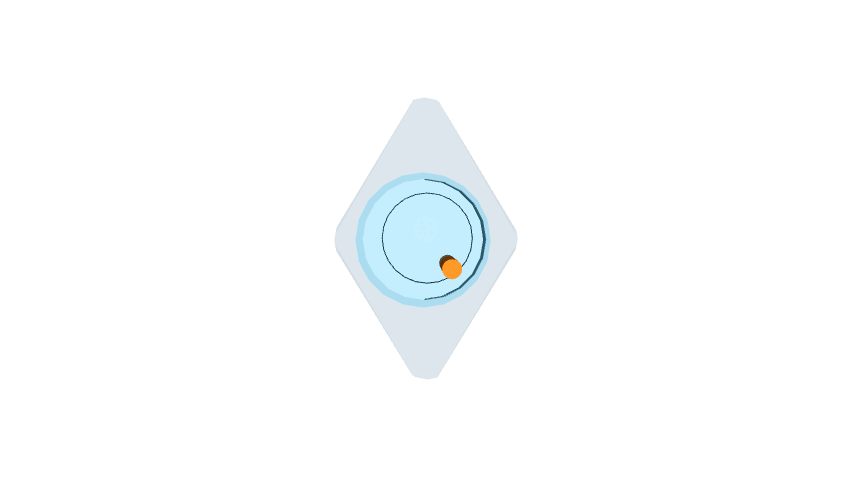
